# The persistence and loss of hard selective sweeps amid admixture in ancient Eurasians

**DOI:** 10.1101/2025.10.14.682443

**Authors:** Mariana Harris, Ziyi Mo, Adam Siepel, Nandita Garud

## Abstract

The extent to which human adaptations have persisted throughout history despite strong eroding demographic events such as admixture, genetic drift, and fluctuations in selection pressures remains unknown. Understanding which adaptations were resilient to such forces may shed light on traits that were important for humans across time. Yet, detecting selection from ancient DNA is challenging due to severe degradation of the data and/or signal. Here we detect selective sweeps using a domain-adaptive neural network (DANN) trained on simulated data and applied to more than 800 ancient and modern Eurasian genomes spanning the last 7000 years. We show that the DANN can account for simulation misspecification, or discrepancies between simulations and real aDNA, improving the ability to detect sweeps in real data compared to standard convolutional neural networks or standard statistics. Application of the DANN to data recovered 16 known sweeps at loci including *LCT*, *HLA*, *KITLG,* and *OCA2/HERC2*, and revealed 32 novel sweeps. All identified sweeps were classified as hard, consistent with historically low population sizes. While some sweeps were lost over time, 14 sweeps at loci involved in functions including neuronal, reproductive, pigmentation, and signaling traits persisted from the earliest to the most recent time periods. In most cases, the most frequent haplotype remained at high frequency across time. Together, these results indicate that hard sweeps predominated in ancient Eurasians and that several ancient selective events were resilient to strong admixture events.

**Significance statement:** The extent to which human adaptations have persisted despite strong eroding forces such as admixture, drift, or fluctuations in selection pressures remains unknown. Understanding which loci are particularly resilient to such forces may shed light on the traits that were important for humans across time. Using a domain-adaptive neural network that accounts for simulations with misspecified demography relative to the data, we discover several sweeps at loci encoding neuronal, reproductive, pigmentation, and signaling traits persisted from the earliest time periods to the present, revealing the resilience of these sweeps to strong admixture events. Moreover, we find that hard sweeps, driven by single beneficial mutations, dominated throughout human history, consistent with the historically low human population sizes.

## 1. Introduction

The growing availability of ancient DNA (aDNA) has revolutionized our ability to study how evolution has shaped human populations over the past ∼12,000 years. The transition from mobile hunter-gatherer groups to sedentary, agriculture-based societies introduced profound selective pressures including shifts in diet, sustained contact with domesticated animals, and heightened pathogen exposure (1, 2). During this same period, repeated waves of migration and admixture among Western Hunter-Gatherers, Anatolian early farmers, and Steppe pastoralists continually reshaped the genetic landscape of Eurasia. These demographic events may have diluted or masked historical selective sweeps in present-day genomes (3), leaving the prevalence, persistence, and modes of selective sweeps across Eurasian populations largely unresolved.

Characterizing the targets as well as the mode and tempo of positive selection in aDNA can reveal the mechanisms and rate of human evolutionary change. However, the ability to detect adaptation in aDNA can be challenging for a number of reasons, including low read coverage, short read lengths, high levels of missing data, and the complex, often poorly characterized demographic history of human populations. Despite these challenges, several studies have shown that directional selection in humans may have been widespread (3–5). However, these studies have largely been powered to detect classic ‘hard’ sweeps, in which a single adaptive variant rising to high frequency leaves behind a characteristic dip in diversity with a single dominant haplotype. In addition to hard sweeps, there may have been ‘soft’ sweeps, whereby multiple adaptive variants rise to high frequency simultaneously given large mutational inputs or abundant standing genetic variation (SGV) at the onset of selection (6, 7). Soft sweeps are more challenging to detect given that they leave behind more subtle signatures in the data due to there being multiple haplotypes rather than a single haplotype at high frequency (8, 9). Given the combination of data challenges, which can generate misleading signals that appear adaptive but actually stem from demographic forces or data artifacts (10–13), and the difficulty in detecting soft sweeps, it remains unknown how many historical sweeps have been missed and whether they were hard or soft.

Deep learning methods have emerged as a powerful tool in population genetics to address a wide variety of inference problems from genomic data including demographic inference (14, 15), estimating recombination rates (16, 17), and detecting selection (16, 18–22). In particular, convolutional neural networks (CNNs) have proven particularly effective in detecting selective sweeps, largely due to their ability to extract complex patterns from noisy, high-dimensional population genetic data. Notably, CNNs can natively handle images of raw genotype matrices as input data, eliminating the need to extract summary statistics (16, 18). Despite their flexibility and strong performance on modern data and other organisms (16, 19), deep learning methods have not yet been applied to aDNA to detect selection, although they have begun to be applied to temporal genomic data (22).

A key limitation to current approaches is that they rely on large amounts of labeled training data, which are typically generated through simulations based on simplified models that are restricted in their ability to fully capture the complexities of real genomic data. Discrepancies between simulated and real datasets can arise from inaccurate assumptions about demography, mutation and recombination rates, or from data artifacts. Additionally, modeling features such as large effective population sizes (*N_e_*), heterogeneous recombination landscapes, or complex demographic histories can be computationally intensive (17, 23), making such simulations impractical at the scale required for deep learning.

This mismatch between simulated training data and real genomic data, known as a simulation misspecification (24), can reduce model accuracy and lead to inferences from data that are not robust. Several strategies have been proposed to address this issue including adaptive re-weighting of training examples (19, 25, 26). Domain adaptive neural networks (DANNs) (27, 28), have recently been proposed as another alternative to mitigate simulation mis-specification (27).

Domain adaptation aims to improve generalization by enabling a model trained on data from a source domain, in this case simulated data, to perform well on a target domain with different properties, such as real population genomic data (29). This technique is widely applied in computer vision: for example, facial recognition models trained on high-quality studio images can be adapted to perform more reliably on lower-quality surveillance footage (30, 31). It is also used in natural language processing, where models trained on reviews of books may require adaptation to accurately interpret sentiments in reviews of other products (32, 33). In biology, domain adaptation has been used to predict transcription factor binding across distinct species (34). Building on these applications, Mo and Siepel demonstrated that domain adaptation could also be leveraged to improve population genetic predictions, including detecting selective sweeps, inferring selection strength, and estimating recombination rates in the face of demographic misspecification (27).

Given the unique challenges of aDNA, domain adaptation presents a powerful framework for characterizing selection across different periods of human history. Here we propose a novel application of a DANN to distinguish between hard sweeps, soft sweeps, and neutrality in aDNA and modern DNA. We find that hard sweeps were more common than soft sweeps throughout human history, consistent with historically low population sizes, and that several sweeps have persisted over multiple time periods, implying resilience to major demographic events and potentially sustained selective pressures over human history.

## 2 Results

### 2.1 Data

In this work, we train a DANN to detect selective sweeps from 708 previously published (35–47) aDNA samples from Europe that we analyzed previously (48), dated between 6500 and 1345 years before present (BP) (**Fig. 1A,B**). These samples are from populations that underwent major admixture events, including the migration of Anatolian farmers into Europe and their admixture with local Mesolithic hunter-gatherers around 8,500 BP, as well as the mixing of European farmers with steppe pastoralists at the onset of the Bronze Age ∼5000 BP (49) (**Fig. 1C)**. This transitional period is particularly important for studying adaptation, as it has been hypothesized that admixture has obscured selective sweep signatures in modern humans and as a result the extent of selection has likely been underestimated (3).

**Figure 1.**
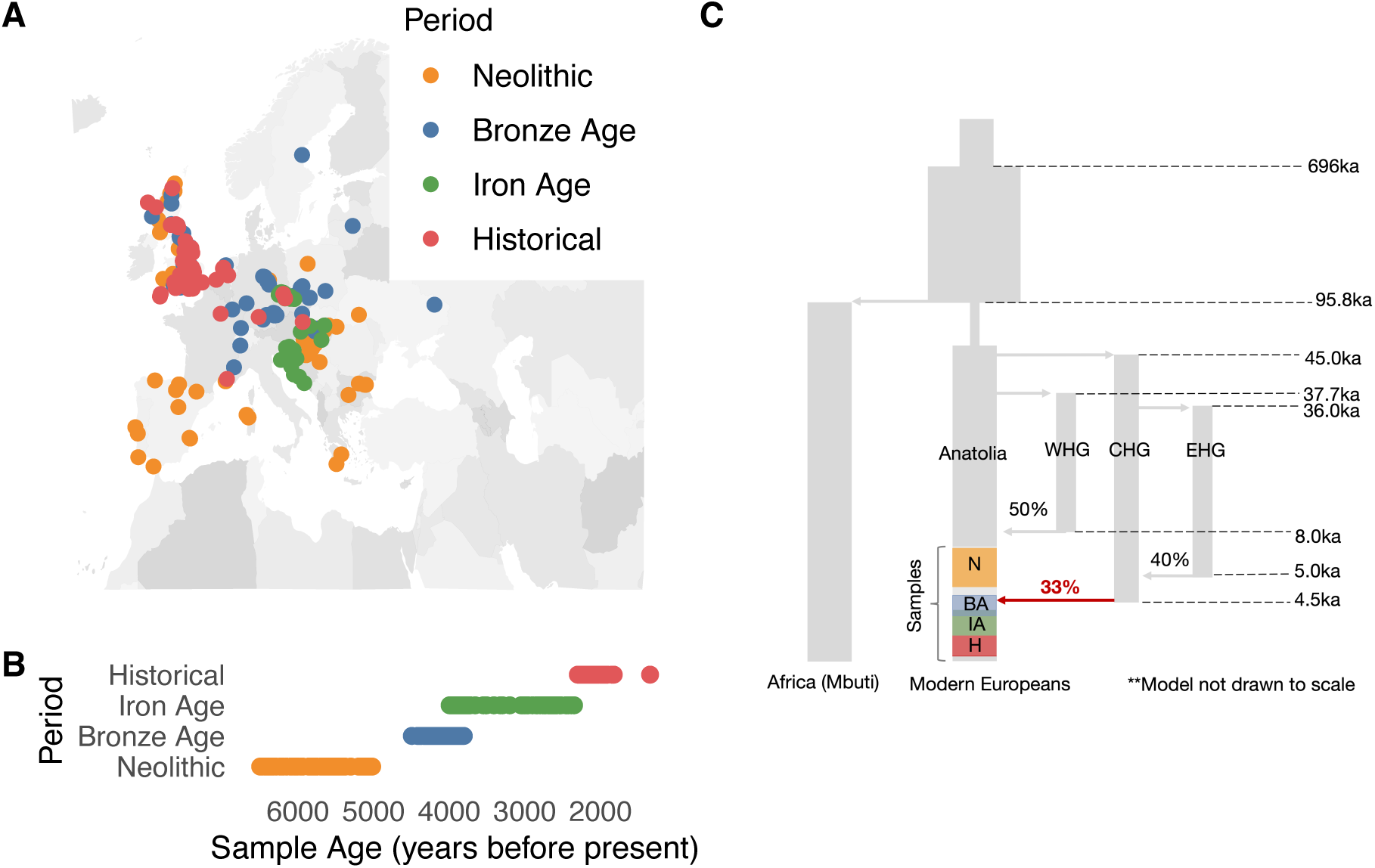
Location and age of samples included in study. **(A)** The locations of the 708 ancient human samples colored by their corresponding time periods. **(B)** Archeological or radiocarbon dates for each sample in years before present (BP). Each data point represents one sample and the colors indicate broader groupings according to four time periods. **(C)** Diagram of West Eurasian population history. Modern Europeans are composed of three main ancestries: Western Hunter-Gatherers, Anatolian early farmers and Steppe pastoralists. The vertical grey segments represent distinct population branches and arrows represent population splits and admixture events. Shown are four main branches: the African (Mbuti) branch, Eurasian branch, West Hunter-Gatherers (WHG), Caucasus Hunter-Gatherers (CHG) and Eastern Hunter-Gatherers (EHG). Highlighted are the time ranges which our samples come from in the Eurasian branch. Also highlighted are the admixture proportions and timing of events. The red arrow depicts a major admixture event overlapping the timeframe during which our samples were collected. The full schematic of the model with all associated parameters can be found in Souilmi et al. 2022 and original studies from which these parameters were estimated (3, 53–55).

Based on direct radiocarbon dates and archaeological context, the samples were grouped into four chronological periods based on *f4* statistics, time period (based on direct radiocarbon dates or precisely dated archeological contexts), and geographic location, resulting in genetic homogeneity within each group with little or no amounts of ancestry from other populations that entered Europe at the time, as described previously (40, 48). The four groups were as follows (**Fig. 1B**):

#### Neolithic (N)

Individuals of European Hunter-Gatherer and Anatolian farmer ancestry dated between 6500 and 5019 BP.

#### Bronze Age (BA)

Individuals from the Bell Beaker cultures of Western and Central Europe, dated between 4495 to 3808 BP.

#### Iron Age (IA)

Individuals from Iron Age Britain and Western Europe dated between 3995 to 2350 BP.

#### Historic period (H)

Individuals from Roman and late antique periods between 2300 to 1345 BP.

To ensure data quality, we only included samples for which a number of criteria could be met (**Text S1**), including requiring hybridization capture on at least 1.2 million positions, having a minimum of 15,000 SNPs such that robust population genetic inferences could be performed, lacking significant contamination on the mtDNA or X chromosome (in males), being unrelated up to the third degree, and being treated with the same Uracil-DNA Glycosylase process during library preparation. The last two bases were trimmed from each read to exclude the most damaged regions of aDNA. After selecting high quality samples, to have similar power to detect sweeps across different time periods, we chose the 177 samples with highest coverage for each time period for further analysis, resulting in a total of 708 genomes (**Table S1**). These 177 samples were subsequently down sampled per analysis window to the 150 samples with the least amount of missing data.

Additionally, we analyzed data from 99 modern European samples (CEU) from the 1000 genomes project (50). We restricted the 1000 genomes samples to the 1.2 million positions that were in the aDNA capture array.

### 2.2 Architecture of the DANN for sweep detection

Since the goal of the DANN is to both classify sweeps from neutrality and unlearn differences between domains, a DANN differs from a more traditional neural network classifier by including not only a classification branch, but also including a discriminator branch that distinguishes between a *source* domain (e.g. simulations) and a *target* domain (e.g. real data) (**Figure 2**). One strategy for domain adaptation, which we use here, is the addition of a gradient reversal layer (GRL) (28). During backpropagation, the sign of the gradient of the loss of the discriminator is reversed through the GRL, penalizing features that discriminate between domains and promoting domain-invariant features that are essential for sweep classification (**Text S2**).

**Figure 2.**
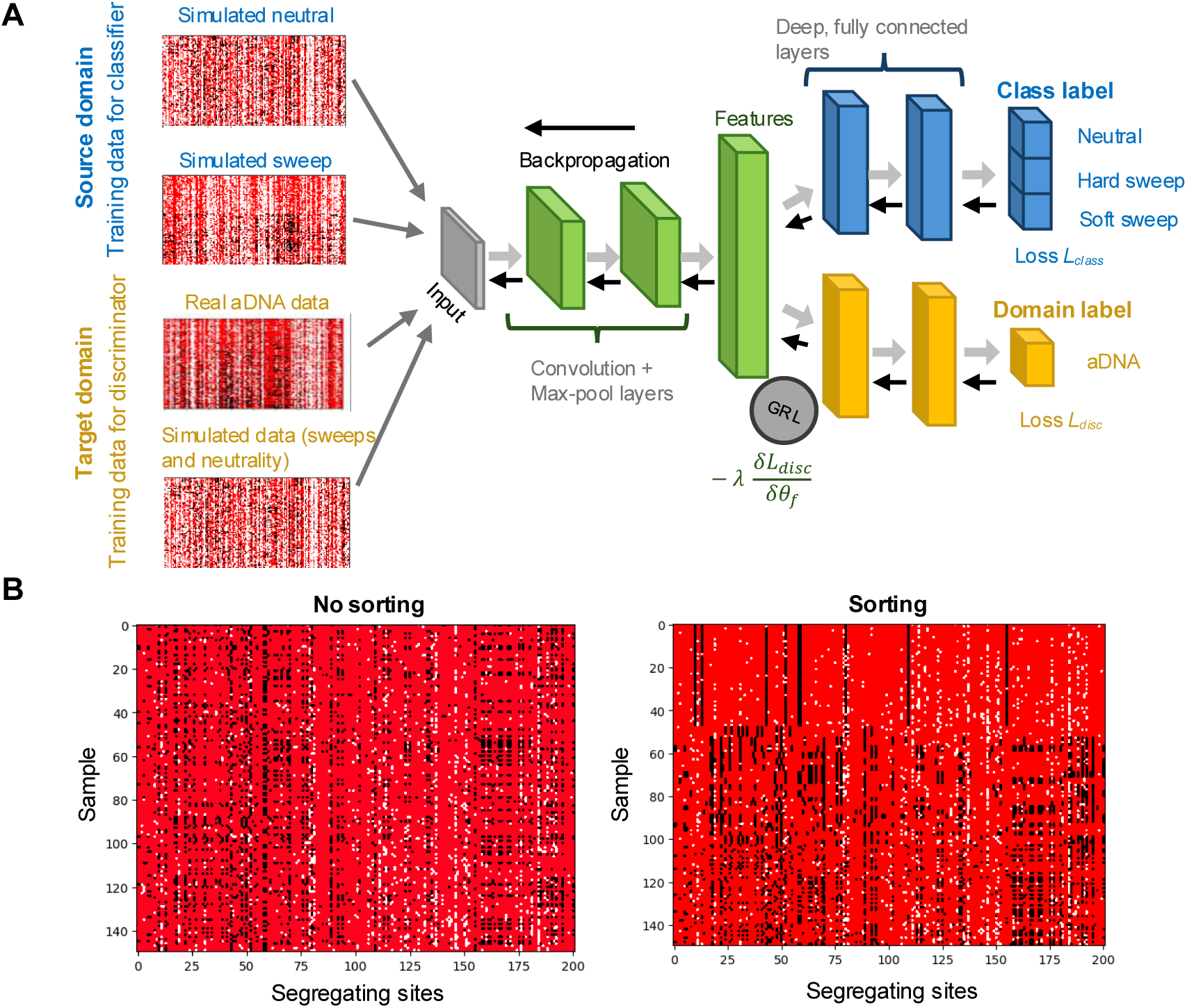
Domain Adaptive Neural Network for detection of selective sweeps. **(A)** DANN Architecture. Haplotype images from both the source and target domains are passed through a series of convolutional layers, with dimensionality reduced by max-pooling steps (green). The output is flattened into a feature vector, which is then processed by two branches of fully connected layers: the classifier, which predicts the sweep class (blue), and the discriminator, which distinguishes between domains (yellow). During backpropagation, the GRL inverts the loss from the discriminator, discouraging the model from differentiating between the domains and promoting domain-invariant features. The grey arrows indicate the forward pass and black arrows indicate backpropagation. **(B)** Genomic data representation. Genomic data is represented as images, with rows corresponding to sampled haplotypes and columns to segregating sites (201 SNPs). Each pixel represents the occurrence of a specific allele with major allele shown in red, minor allele in black, and missing data in white, coded as -1,1 and 0 respectively. On the left, we show a haplotype image of a simulated partial hard sweep where no sorting was applied. To the right we show the same image but sorting the rows (haplotypes) by frequency.

As input to the model, we provided images of haplotypes sorted by the frequency of most to least common haplotype. Because of the low coverage nature of aDNA as well as ascertainment bias in calling ancient SNPs, previous work has shown that heterozygous sites are not always reliably determined. To address this issue, we used our previous approach (48) to ‘pseudo-haplodize’ the data by randomly selecting one of the reads mapping to a position and assigning the genotypes of the read as the genotype of the sample at that site (**Text S2**). This data was then used as input to the DANN.

The images provided to the DANN are *n×S* bi-allelic genotypic matrices representing the allelic states at *S*=201 segregating sites across *n*=150 pseudo-haplotypes (**Figure 2B**). These parameters reflect our previous findings (48) that a window size of 201 segregating sites spanning approximately ∼450 kb in aDNA is adequate for capturing signals of selection in this dataset using haplotype homozygosity statistics. In addition, sorting haplotype images based on haplotype distances or frequencies has been shown to be essential for strong performance of CNN models (16, 18, 51, 52). We tested the ability of four different sorting approaches to distinguish hard sweeps, soft sweeps, and neutrality (**Fig. S1, Text S3**): (1) sorting by distance to the most common haplotype, designed to emphasize elevated haplotype homozygosity typical of hard sweeps, (2) sorting haplotypes by highest to lowest frequency, designed to emphasize the presence of multiple high-frequency haplotypes characteristic of soft sweeps, (3) sorting a central 51 SNP window by the distance to the most common haplotype within a 201 SNP window, designed to allow the model to utilize information both at the sweep center and flanking regions, and (4) sorting a central 51 SNP by haplotype frequency within a 201 SNP window. While all sorting approaches had improved power relative to no sorting, sorting by haplotype frequency had the highest power for detecting soft sweeps, as this sorting strategy likely best presents to the model the presence of multiple haplotypes at high frequency characteristic of soft sweeps. Thus, in all analyses going forward, we sort by haplotype frequency. In the future, however, leveraging permutation-invariant layers may improve the generalizability of our model for other population genetic tasks (52).

### 2.3 Benchmarking DANN performance

#### 2.3.1 Robustness to domain shift and missing data

Before applying a DANN to aDNA, we first assessed its ability to distinguish neutrality, hard sweeps, and soft sweeps under varying levels of misspecification between the source and target domains. Specifically, we simulated different degrees of mismatch in demography and missing data rates between the target and source domains, as these two factors were expected to generate the greatest discrepancies between real data and simulations. In all scenarios, the target domain, used as a proxy for real aDNA, consisted of simulations under a previously inferred admixture model describing ancient Europeans (3, 53–55) (**Fig. 1C**) with 43% missing data per base pair, matching the missing data rate of the aDNA samples analyzed in this study (**Text S4, Fig. S2**). The source domain varied in both demographic history and missing data rate, with different variants of the admixture model (**Fig. S3**) and a constant *N_e_* = 10^4^ model, and with either low (5%) or high (43%) missing data.

To evaluate the effect of the discriminator branch on the ability of the DANN to correctly classify sweeps, we compared the performance of the DANN to that of an otherwise identical CNN without a discriminator in two evaluation settings. In the hypothetical best-case scenario, the source and target domains share the same demographic model and missing data rate, such that there is effectively no domain shift. In this setting, the CNN is trained and tested on data drawn from the same distribution, and the gradient reversal layer of the DANN is therefore expected provide no benefit. This setting serves as an upper baseline on classification performance. By contrast, in the mismatched setting, the same CNN architecture is trained on the source domain but evaluated on a target dataset that differs in demography and/or missing data patterns. This scenario reflects a typical application of a CNN, in which the distribution of the target data is unknown and shifted relative to the training distribution (standard CNN). Comparing CNN performance in the matched and mismatched settings to that of the DANN allows us to quantify the extent to which the discriminator branch improves classification performance specifically under domain misspecification.

We found that for all simulation scenarios, the DANN outperformed the CNN, albeit modestly, when the target and source domains did not match, demonstrating its ability to mitigate misspecification (**Fig. 3, Fig. S4, Fig. S5**). While there is improvement, the DANN cannot completely correct for misspecification, as the hypothetical best-case scenario with matching domains always performs best. Additionally, we found that the DANN could correct for mismatch in demography better than mismatch in missing data rates between target and source domains with the DANN performing slightly better in mitigating the most extreme misspecification in demography (constant *N_e_* vs. Admixture; area under the precision-recall curve (AUPRC) of 0.814) compared to the most extreme misspecification in missing data rates (5% vs 43%; AUPRC =0.80) (**Fig. 3A**). Likely, the lower power of the DANN under the missing data misspecification scenario reflects the severity of the mismatch in domains. Increasing the levels of missing data directly removes information and reduces the signal available for classification. By contrast, demographic misspecification, while shifting patterns of variation, still preserves key sweep-associated features, allowing the DANN to better learn representations that are invariant to demographic differences.

**Figure 3.**
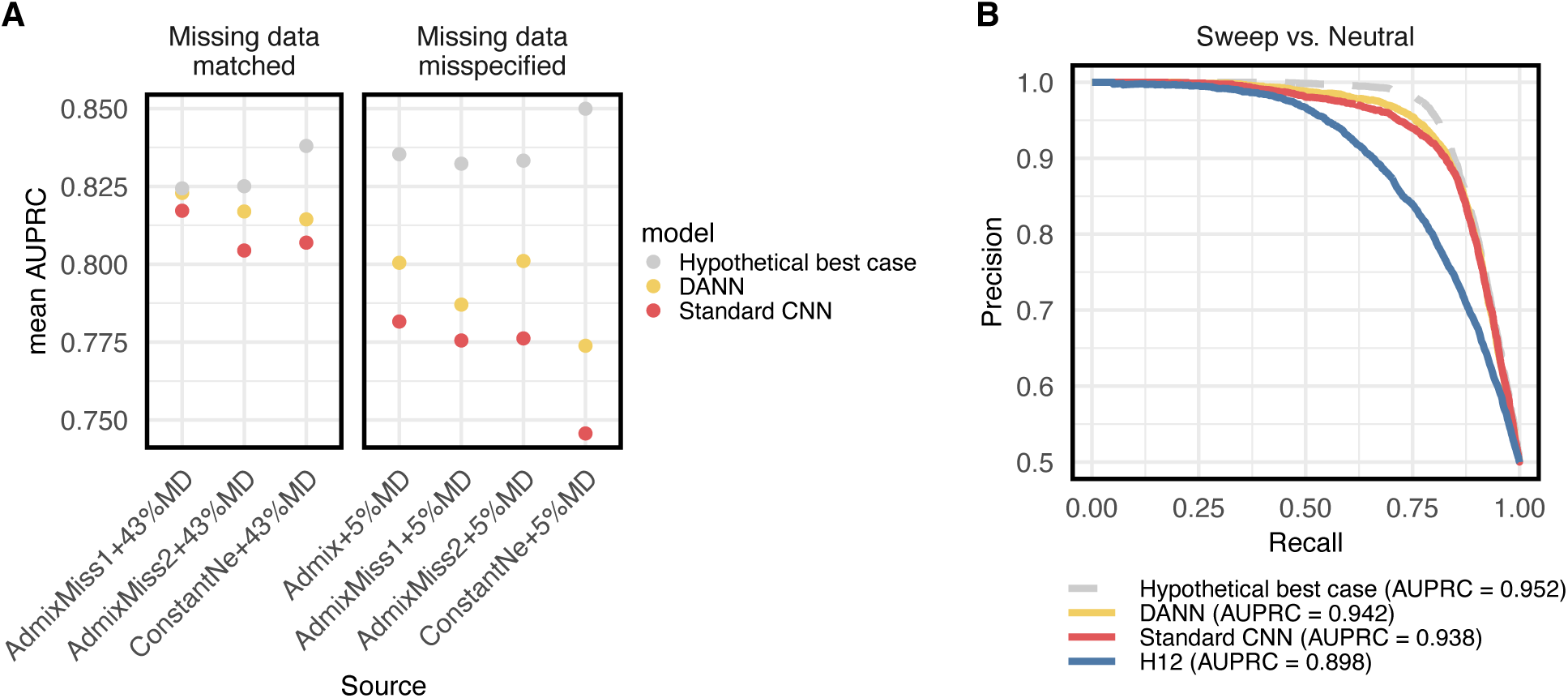
Domain adaptive neural network improves the detection of selective sweeps in simulated data that mimics aDNA. **(A)** Ability of the DANN to correctly classify hard sweeps, soft sweeps, and neutrality. In all scenarios, the target domain, used as a proxy for real aDNA, is composed of simulations of the human admixture model (3, 53–55) with a 43% missing data (MD) rate per base pair. The source domain is specified on the x-axis, where “AdmixMiss1” and “AdmixMiss2” refer to models drawn in **Fig. S3**. In red and yellow are a CNN and DANN, respectively, tested on mismatching domains with the source having different demographic models and/or missing data rates from the target. This is compared with a hypothetical best case simulation benchmark CNN (grey), representing an upper bound on performance. This benchmark was trained and tested on matching source and target domains. **(B)** Precision-recall curve for detection of sweeps (hard and soft) vs neutrality using a constant *N_e_* = 10^4^ model as a source domain and the human admixture model as a target domain, both with an average of 43% missing data per site.

#### 2.3.2 Generalization to aDNA

Since aDNA will ultimately be the target data for the DANN, to test its ability to discriminate between neutrality, hard, and soft sweeps in this more realistic scenario, we trained the DANN with real aDNA for the target domain and simulated data for the source domain. We found that when we evaluated the performance of the DANN in this scenario, the performance was similar to that of the scenarios where both target and source were simulated data (**Text S5, Fig. S6)**.

#### 2.3.3 Ability of the DANN to detect sweeps of varying strengths, frequencies, and softness

The ability to detect selective sweeps may vary with the strength of selection, whether the sweep is partial or complete, and the softness of the sweep. To evaluate the performance of the DANN across evolutionary scenarios, we varied selection strength, sweep frequency, and sweep softness in simulations where the source domain was the constant *Ne* model and the target domain was the human admixture model. We simultaneously compared the performance of the DANN to that of H12, a haplotype-based statistic capable of detecting both hard and soft selective sweeps (56) that was recently applied to aDNA (48).

Specifically, we tested the performance of the model on weak (s∼[0.005,0.05]) and strong (s∼[0.05,0.1]) selection. We found that when selection is weak the DANN is able to distinguish sweeps from neutrality with an AUPRC of 0.89, and when selection is strong, the AUPRC is 0.98 (**Fig. S7A**), outperforming H12 (AUPRCs of 0.827 and 0.977 for weak and strong selection, respectively). This was also the case when evaluating the ability of detecting hard sweeps and soft sweeps separately (**Fig. S7, Fig. S8**). Additionally, we tested the performance of the model on partial and complete sweeps. We found that the DANN outperforms H12 in all scenarios tested with the exception of complete hard sweeps with strong selection (**Fig. S7**).

Finally, we tested the ability of the DANN to detect soft sweeps of varying softness modulated by rate of input of adaptive mutations θ_A_ = 4*N_e_*µ_A_, where µ_A_ is the adaptive mutation rate. We asked how often soft sweeps are misclassified as hard sweeps and vice versa, conditional on being distinguishable from neutrality (**Fig. S8)**. We found that the majority of hard sweeps (82%) are correctly classified as hard, with this percentage increasing to 93% for strong selection. Similarly, we found that the majority of soft sweeps (∼75%) are correctly classified as soft, with this percentage increasing to 86% as θ_A_ increases to 5 when the signatures of hard vs soft sweeps become most distinct.

### 2.4 Application of the DANN to aDNA and modern humans

Having confirmed the ability of the DANN to detect selective sweeps in simulations, we now apply the model to aDNA to discover selective sweeps. We focus on a DANN trained using aDNA as the target domain and simulations generated under the admixture model shown in Figure 1 with a 43% missing data rate as the source domain (section 2.3.2). We applied the DANN genome-wide in sliding windows of 201 SNPs, advancing each window by 10 SNPs. To ensure predictions are well supported by more than one window, we averaged the predicted probabilities every five consecutive, overlapping windows, generating the final class predictions (**Text S6**). The class (hard sweep, soft sweep, or neutral) with the highest mean predicted probability across the five windows was then assigned to the genomic position corresponding to the location of the central window of the 5 consecutive windows. We show the resulting scan across the aDNA time transect in **Fig. 4A**, where windows predicted to be hard sweeps are shown in red, soft in blue, and neutral regions in grey. In addition, In **Fig. 4B**, we show a scan on the modern European population. This scan was generated using a different model trained on modern human data as the target domain (**Text S5**).

**Figure 4.**
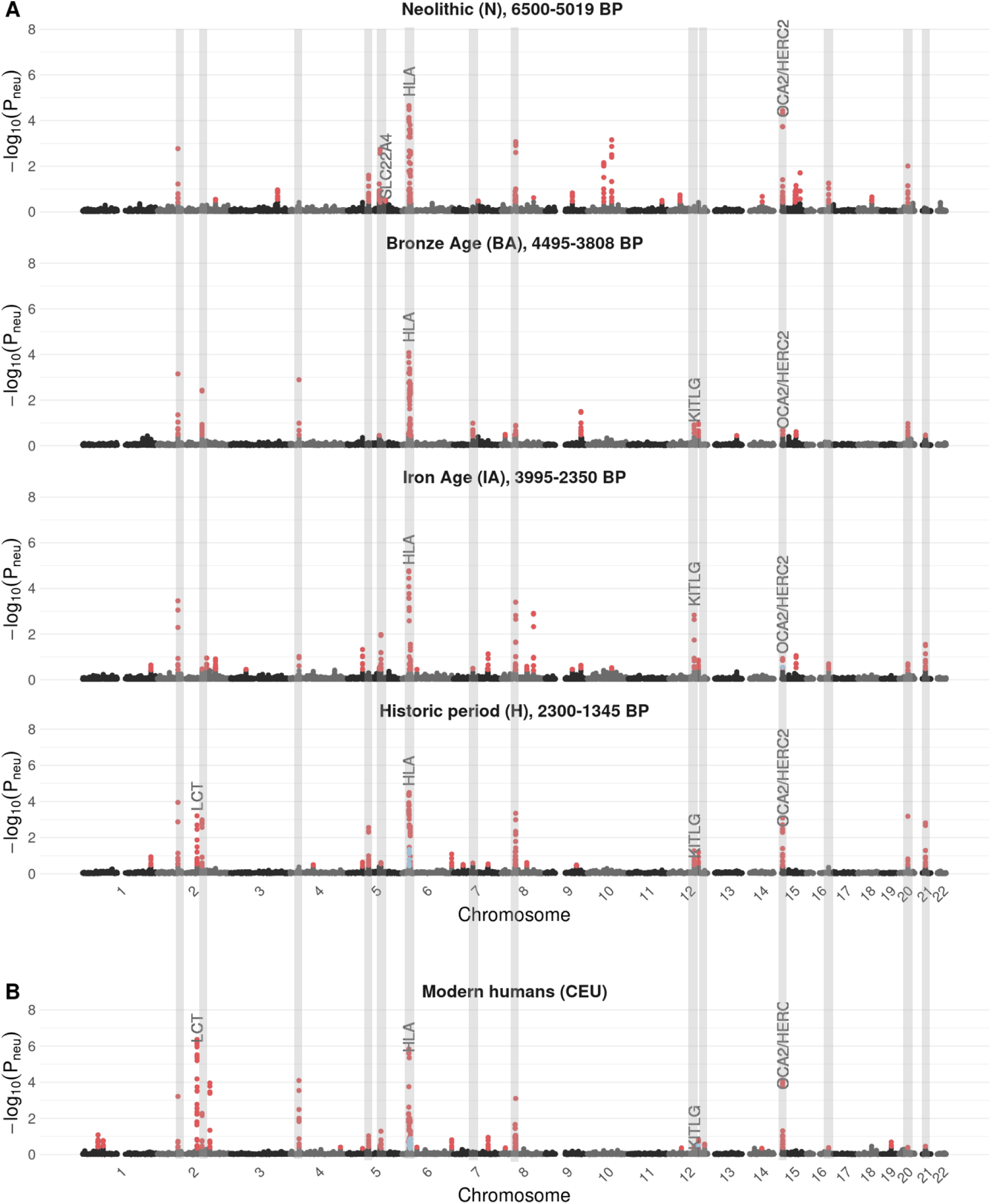
Genome-wide selection scan with the DANN in ancient and modern data. Results for the DANN are shown for all four ancient time periods **(A)** and for modern Europeans **(B)**, highlighted at the top of each panel. The time period range is specified as years before present (BP). The y-axis shows the probability of selection -log(P_neu_) predicted using the DANN, whereby a high value indicates the window is likely under selection. The x-axis shows the genomic position. The DANN was trained using a human admixture model with missing data as the source domain and aDNA data as target domain. Windows predicted as hard sweeps are colored in red and windows predicted as soft are colored in blue. Canonical sweeps previously reported in the literature are indicated above their respective genomic positions. We highlight with grey vertical bands 14 sweeps that are present across the two ends of the major admixture event that occurred ∼4.5 kya, that is sweeps that are detected in earlier periods (N or BA) and are also detected in later periods (H or CEU).

To avoid calling the same selective sweep multiple times, we grouped consecutive non-neutral windows into a single sweep. Additionally, to ensure distinct selective events, we further required distinct sweeps to be at least 1.5 Mb apart. Within each sweep, we identified the representative window as the one with the highest sweep probability (the highest –log(probability of neutrality)).

We identified a total of 48 unique sweeps in ancient humans (**Fig 4A**) and 28 selective sweeps in modern humans. Among the 48 ancient human sweeps, over half the sweeps recurred across multiple periods with 5 detected in all periods, 8 in three, 12 in two, and 23 in only one (**Fig. S9, Table S2**). Additionally, 18 ancient sweeps were found to persist in modern humans, resulting in 58 unique sweeps across all ancient and modern periods. In Fig. 4, grey vertical bands highlight 14 sweeps that span the major admixture event that occurred ∼4.5 kya, being detected in earlier periods (N or BA) and persisting into later periods (H or CEU).

To assess the robustness of our sweep inferences in ancient humans, we trained two additional models varying the source domain, including a constant *Ne* instead of admixture model with 5% instead of 43% missing data rates, and a model trained on soft sweeps from SGV instead of *de novo* mutations. We observed high concordance across models both in sweep detection and classification, with 44 sweeps overlapping in all three scans (**Fig. S10A**, **Fig. S11, Table S2**).

Out of the 48 sweeps detected in ancient humans (**Figure 4A)**, 16 overlap with recent studies (3–5, 35, 48, 57, 58). We do not consider sweeps as novel candidates if they are within 1Mb away from genes that have been previously identified as sweep candidates. Among the 16 previously discovered sweeps are 5 well known targets of selection, as highlighted in **Fig. 4**. The strongest signal across all time periods corresponds to the *HLA* region and neighboring gene *ZKSCAN3*. *HLA* encodes cell surface proteins that are involved in the adaptive immune system and has long been recognized as a target of selection (59, 60) and *ZKSCAN3* is involved in transcriptional regulation of autophagy-related genes and was reported as under selection in Mathieson et al. 2015. We also detect a sweep spanning the *OCA2/HERC2* genes, which is associated with light eye color in Europeans (35). We identify this signature across all four time periods, which was not observed in earlier scans of the same time transect (48, 58). Our scan also recovers a strong sweep signal at the *LCT* locus, which is associated with lactase persistence into adulthood (61, 62). This signal is restricted to the Holocene (H) period and the CEU population (**Fig. 4**), consistent with the rapid rise in frequency of the causal variant rs4988235 during this time. Notably, this allele was absent in Europe prior to the arrival of Steppe pastoralists in the Bronze Age and therefore could not have been under selection earlier (35, 48, 58, 63–65). We also highlight a sweep overlapping the gene *KITLG*, associated with light hair and skin pigmentation and recently identified as a target of selection, and *SLC22A4,* associated with Crohn’s disease (4). We summarize the number of sweeps that overlap with previous work in **Table S3** and **Fig. S12,S13**.

The remaining 32 ancient sweeps we identify represent novel candidates that, to our knowledge, have not been identified in previous studies. To understand if the genes within these 32 sweeps are enriched for any functions, we annotated all protein coding genes within 300 Kb distance upstream and downstream of the central SNP in the window with highest probability within a sweep using Ensembl Variant Effect Predictor (VEP) (**Supplementary Table S4**). We next performed an enrichment analysis for previously identified genome-wide association study (GWAS) annotations on the set of mapped genes using Functional Mapping and Annotation of Genome-Wide Association Studies (FUMA) (66). We observed an enrichment in GWAS categories for human traits, including body measurements (e.g., hip circumference), neurological traits (e.g., brain morphology), metabolic traits (e.g., triglyceride levels), and immune- or autoimmune-related traits (e.g., type 1 diabetes, asthma). These results are reported in **Fig. S14.**

### 2.5 Hard sweeps were more common than soft sweeps in ancient and modern humans

We next analyzed whether sweeps discovered in aDNA and modern humans were predominantly hard or soft. The DANN classified all 58 sweeps (ancient and modern) in **Fig. 4** as hard. In three cases, we observe that windows on the edges of the sweep are classified as soft (chromosomes 6 and 15, **Figs. 4 and S15)**. This pattern is consistent with the “soft shoulder effect” (67), where recombination causes regions flanking a hard sweep to exhibit patterns that resemble a soft sweep. However, based on our sweep calling approach, these flanking regions are not classified as independent soft sweeps. Instead, they are grouped with adjacent windows into a single sweep, which is then labeled according to the window with the highest sweep probability.

To evaluate whether the predicted hard sweeps may reflect a high rate of soft-to-hard sweep misclassification by the model, we use simulated data from the human admixture model to estimate the proportion of predicted hard sweeps that may correspond to soft sweeps misclassified as hard. Applying these rates (**Fig. S16**) to our ancient hard sweep predictions, we estimate that, conservatively, 83% of detected sweeps are hard and 15% are soft, indicating that despite any potential for misclassification, hard sweeps were likely more common than soft sweeps in ancient human populations. It is important to note, however, that these error rate estimates are rough approximations based on simulated data, as the true labels in real aDNA are unknown.

Additionally, to assess robustness of our results to the underlying model of soft sweeps, we trained a model in which soft sweeps were simulated from SGV rather than recurrent *de novo* mutations (**Fig. S10B**). We identified a total of 53 unique sweeps in aDNA, 43 of which overlap with the scan trained on *de novo* soft sweeps. Of the 53 sweeps, 38 are classified as hard in every time period in which they appear, 9 are classified as soft, and 6 change from hard to soft over time. However, the predicted probabilities for soft sweeps are low (0.34–0.44), only slightly above the value expected under random assignment among hard, soft, and neutral classes (0.33), indicating weak support for these soft sweeps. By contrast, predicted probabilities for hard sweeps range from 0.34–0.93, indicating much stronger support. Given uncertainty among the sweeps with lower probabilities, we tested a more stringent classification threshold for sweeps by doubling the 0.33 baseline to 0.66. Conditioning on hard and soft sweeps with predicted probabilities greater than 0.66, we identified 15 hard sweeps and zero soft sweeps.

To further validate whether the sweeps detected by the DANN are in fact hard sweeps, we visualized the haplotype structure of the window with the highest probability supporting the sweep classification of all sweeps detected in **Fig. 4**. By visual inspection, in all cases, we observe a single haplotype at intermediate frequency across all windows, consistent with partial hard sweeps, as highlighted in 24 examples in **Fig. S17**. These windows are distinct from soft shoulders found on chromosome 6 of period H (**Fig. S18**), whereby multiple haplotypes are at high frequency, and distinct from windows classified as neutral, whereby no haplotypes are at high frequency (**Fig S18**).

In addition to the above, to further confirm inferences about the softness of sweeps made by the DANN, we compute the value of the haplotype homozygosity statistics H12 and H2/H1 (56) for the sweep candidates detected in Figure 4A. These statistics have been previously used to detect and classify hard and soft selective sweeps, where H2/H1 is expected to be small for hard sweeps and large for soft sweeps, conditional on H12 being larger than expected under neutrality (56). To analyze the sweeps least likely to have H12 values consistent with neutrality, we focused on 23 sweeps in total that had an H12 value greater than the 95th percentile of H12 values from 400,000 neutral simulations under the human admixture model. Using an approximate Bayesian computation (ABC) approach to estimate whether each (H12, H2/H1) pair observed in the aDNA is more likely to arise under a hard or soft sweep model (**Fig. S19, Text S7**), we find that 22 sweeps are better supported by a hard sweep model (BF<1) with 15 showing strong support (BF<0.5). Although this is confirmatory of the most extreme outlier selective sweeps, the DANN also identifies more than 25 additional sweeps whose signatures are indistinguishable from neutrality by H12, demonstrating that the DANN and H12 capture non-redundant signals.

### 2.6 Selective sweeps through time

The massive admixture events in which large fractions of the European population experienced genetic turnover may have important implications for the ability to detect selective sweeps over time. Previous work has suggested that sweeps in ancient populations may no longer be detectable in modern time periods due to a masking effect of admixture and drift (3, 48). Of relevance to our dataset, 33% of the ancestry of the Anatolian branch, from which our samples descend, is inferred to have been replaced by a Caucasus Hunter-Gatherers (CHG) population ∼4.5kya, overlapping samples from the BA and IA periods (**Fig. 1B, C**). We asked whether sweeps detected before this admixture event are detectable subsequently, as this would imply resilience to major events and shared selective pressures through time.

To investigate the potential impact of admixture on sweep detection across time, we quantified the overlap in selective sweep signals across the five time periods studied here. We found that among the ancient time periods (N-H), 35 sweeps are detectable in only one or two periods, suggesting either admixture and drift may in fact have had an impact on sweep detection or there may have been a relaxation in selection pressures (Table S5). However, at the same time, the remaining 13 ancient sweeps are detectable in 3 or more time periods, with 10 persisting into the modern time period.

We asked whether lack of overlap of sweep signals is correlated with the admixture event 4.5kya. To do so, we measured the overlap in sweeps between pairs of ancestral time periods (N-H) using the Jaccard similarity index (*J*), which measures the proportion of shared elements between two sets relative to their union (**Text S8**). We asked whether these observed Jaccard values were larger or smaller than expected under a null where sweeps are randomly distributed across time periods. We found that the farthest time periods, N and H, which have the admixture event separating them, had significantly less sharing of sweeps than expected by chance (*J=0.17,* p-val=0.03, permutation test, **Fig. S20F, Text S8**). This is consistent with some sweeps detected in the earliest time period (N) being lost due to the impact of admixture and/or drift over long timescales. However, the absence of sweeps in later periods may, in some cases, reflect limited power rather than true biological loss. Additionally, we found that two sets of consecutive time periods, (BA, IA) and (IA and H), have higher than expected sweep sharing (*J=0.44-0.45,* p-val <0.05, permutation test, **Fig. S20B-C**, **Text S8**), indicating that on shorter time scales sweeps can persist.

We further investigated whether lack of sweep persistence between time periods could be due to either the effects of admixture or drift, or, due to relaxed selection pressures. To do so, we asked whether the most frequent haplotype, presumably carrying the adaptive mutation, of the 35 sweeps that are present in only 1 or 2 time periods, is detectable in other periods (**Table S6)**. Since haplotypes can mutate over time, we assessed whether the most frequent haplotype at these sweeps is more similar to any haplotype present in subsequent time periods than expected based on the average divergence between random pairs of haplotypes (**Fig. S21, Fig. 5**). In 18 cases the most frequent haplotype was eventually lost, suggesting that admixture and drift may have eroded these sweeps (**Fig. 5A**). However, in 17 out of 35 sweeps, the haplotype can be found in all subsequent time periods, despite the sweep not being detected, suggesting that admixture and drift did not completely erode the presence of the adaptive material over time and that instead selection pressures at these loci may have fluctuated.

**Figure 5.**
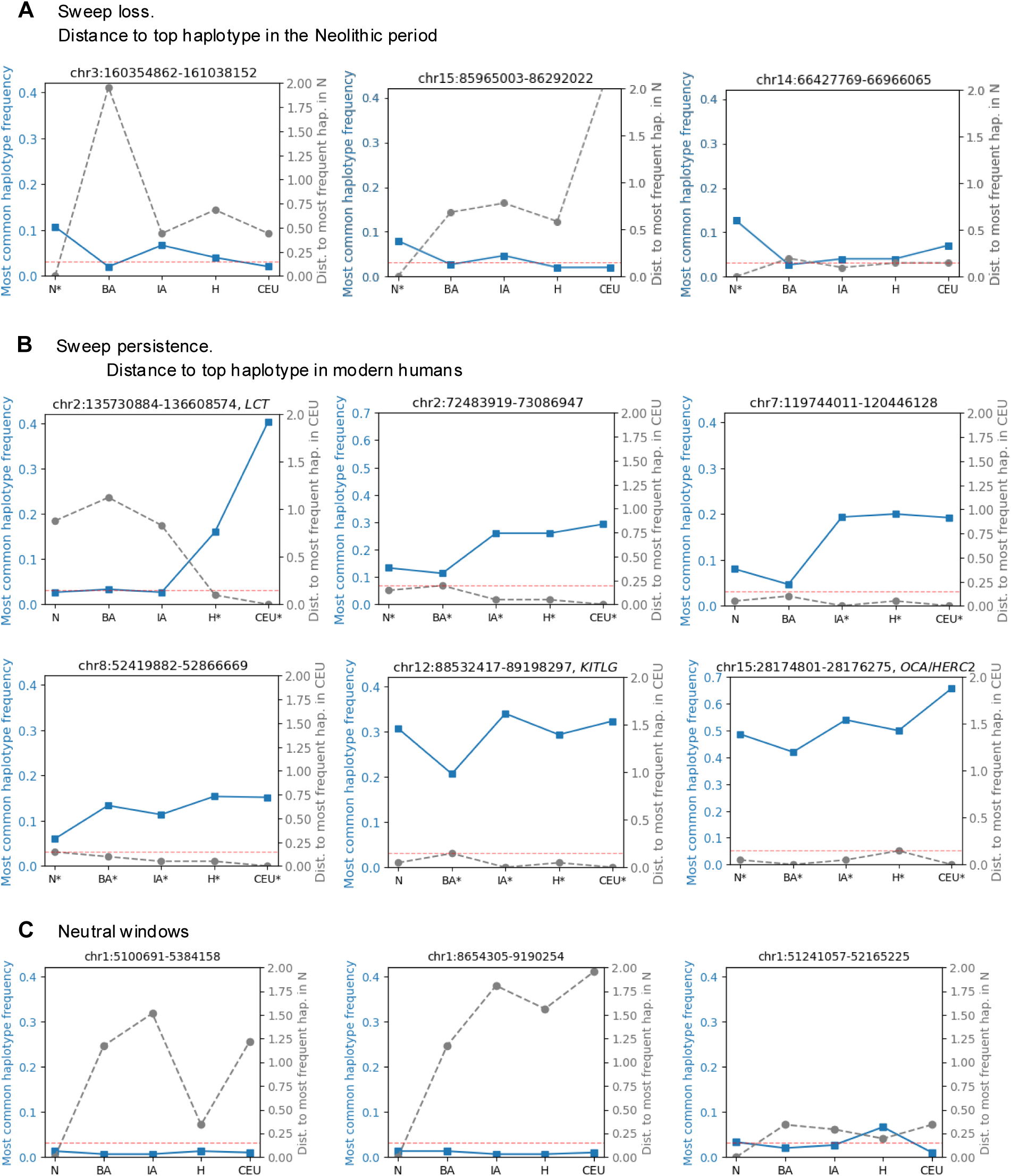
Persistence of the most frequent haplotype across time. **(A)** Three examples of sweeps that are no longer detected with the DANN after time period N. In the two leftmost panels the top haplotype is completely lost, whereas in the rightmost panel the haplotype persists at lower frequency in subsequent periods but is no longer classified as a sweep by the DANN. **(B)** Six examples where the most frequent haplotype persists across multiple time periods. **(C)** Three examples of windows classified as neutral in all time periods. In all panels, the blue line (left y-axis) shows the frequency of the most frequent haplotype in four ancient populations (N–H) and in the modern population (CEU). The grey line (right y-axis) shows the normalized Hamming distance between the most frequent haplotype in each time period and the most frequent haplotype in CEU in the case of **(A)** or in period N in the case of **(B)**, **(C)**. The horizontal red dashed line marks the 1st percentile of the distance between random haplotype pairs, with values above this threshold indicating distinct haplotypes. We highlight with a star (*) the time periods for which the window was

More broadly, despite there being significant non-overlap in sweeps pre- vs post-admixture, 14 sweeps that were identified before the admixture event in either the Neolithic or Bronze Age time periods were also present after the admixture event in the Historical time period and modern humans (highlighted in grey vertical bands in **Fig. 4** and also summarized in **Fig. S9**). This suggests that admixture events may not have obscured all ancient sweeps and that some sweeps may have been subject to sustained selection pressures over time, implying their potential functional importance for human evolution.

We asked whether sweeps that are shared across multiple time periods persist because the same haplotype carrying the adaptive allele remained under selection, or whether the original haplotype was replaced by a distinct haplotype, potentially introduced through admixture. Once again, we calculated whether the most frequent haplotypes of sweeps shared across time periods are more similar than expected. Out of the 14 sweeps detected on either end of the major admixture event spanning our dataset (**Fig. 4**), 9 of these share the same most frequent haplotype across at least 4 time periods, including periods where our method does not detect the sweep. This implies that the sweeping haplotype frequently persisted despite widespread admixture. In **Fig. 5B**, we highlight 6 examples of these sweeps. Among these we include a more recent sweep at the *LCT* locus where the recent rise of the adaptive haplotype is particularly evident. We note that the temporal patterns highlighted in **Fig. 5A,B**, which track haplotype frequencies at selective sweeps, are strikingly different from those in regions classified as neutral by the DANN (**Fig. 5C**), where no haplotype reaches high frequency and the distances between the most frequent haplotypes fluctuate.

The genes identified in the 14 selective sweeps persisting despite the major admixture event fall into a few functional categories: These include neural and cognitive functions encoded by *AUTS2*, *ASCL1*, and *SEMA6A*, of which *AUTS2* was previously discovered to putatively be under selection (68), neuronal signaling and calcium channels encoded by *CACNB4*, exocytosis encoded by *EXOC6B* (*69*), and previously discovered adaptations at pigmentation genes *OCA2*, *HERC2*, and *KITLG* (4, 35). Most of these genes are either found solo within the coordinates of their respective selective sweeps, or with few other genes, narrowing the targets of selection. Contained in sweeps with more genes are metabolic and nutrient processing genes like *PAH* and *SLC38A9*, reproductive and germ cell genes such as *DDX4*, *SPAG4*, and protein quality control and signaling genes like *LTN1*, *USP16*, *CCT8*, and *MAP3K7CL* (**Table S5**). Genes overlapping persistent selective sweeps are not significantly longer than genes genome-wide, suggesting that sweep detection is not biased toward long genes (**Fig. S22**, p-val=0.22). Together, the gene categories present in the 14 sweeps persisting through history highlight functional classes, particularly cognitive and pigmentation, that were potentially of great importance throughout the past 7000 years of history. Future work, however, is needed to fully understand the nature of positive selection at these loci.

## 3. Discussion

In this study, our goal was to characterize the targets and mode of adaptation in ancient humans. To do this, we implemented a domain adaptive neural network that is able to detect and classify selective sweeps in aDNA data and is robust to model misspecification. We applied our model to aDNA from 708 samples spanning ∼7000 years in the past as well as 99 modern Europeans, and identified 48 unique ancient hard selective sweeps and 28 modern hard sweeps, recovering both previously known and novel candidates of selection. Finally, we found that while some sweeps identified in the Neolithic are no longer detected in more recent time periods, 14 sweeps overlapping genes involved in neuronal, reproductive, pigmentation, and signaling traits can be found in the earliest and latest time periods despite the impacts of drift and admixture.

Past studies on humans have generally found few clear examples of hard sweeps in modern genomes (70, 71). Souilmi et al. proposed that this paucity of classic sweeps reflects the impact of complex population history, in particular the masking of sweep signatures by strong admixture events. Using SweepFinder2 (72), which is primarily powered to detect completed hard sweeps, they identified 57 sweeps across ancient Eurasian human genomes, including many sweeps that are undetectable in modern data. Our DANN recovers several sweeps that show a similar pattern: of the 24 sweeps detected in the most ancient time period of our study, the Neolithic, 41% are not detected in the following periods (**Fig. S9**). However, our results also reveal additional dynamics not previously captured: the persistence of sweeps across time periods, even those most impacted by admixture. First, we observed sweeps in which the most frequent haplotype is shared across multiple time periods, even when the sweep is not detected by the DANN in all time intervals (**Fig 5**). This pattern might reflect standing genetic variation, where the haplotype has not been driven by selection to a high enough frequency to be detected by the DANN. We also detect cases where the most frequent haplotype appears in non-consecutive periods, suggesting more complex evolutionary dynamics such as fluctuating selection or a temporary replacement of the most frequent haplotype driven by admixture or migration that obscure detection of the sweep in some time intervals.

The finding that hard sweeps were more common than soft sweeps in aDNA suggests that adaptation in humans has largely been mutation limited, or that SGV available to seed adaptation was low. Even accounting for a 16% misclassification rate of soft sweeps as hard sweeps (**Fig. S16**), hard sweeps remain the dominant mode of selection in aDNA. The high frequency of hard sweeps in the data is consistent with the notion that hard sweeps are expected to dominate in populations with small population sizes, where the input of new adaptive mutations is low (7, 73). In ancient human populations, the effective population size is estimated to be relatively small (55) ( *N_e_*∼10^4^), therefore the input of new adaptive mutations was likely moderate (*Θ*_A_ ∼ 0.001), making adaptation through hard sweeps likely. We confirmed the DANN is not inherently biased toward hard sweeps by testing it on a North American population of *Drosophila melanogaster* where there are three well-characterized cases of soft sweeps (74–80) and successfully recovered all known sweeps as soft (**Text S9, Fig. S23**). Nonetheless, given the inherent challenges of distinguishing between hard and soft sweeps, especially in regions of the parameter space where their genomic signatures overlap, additional work will be needed to fully resolve the rapidity of human adaptation.

Domain adaptation provides meaningful improvement on supervised machine learning methods for analyzing genomic data. With the significant increase in available aDNA samples over the past decade (81, 82), domain adaptation may be especially valuable, as aDNA analyses may be susceptible to false positives due to unmodeled or unknown demographic events as well as the overall poor quality of the data. Domain adaptation was recently applied to SIA, a method that detects selection using the ancestral recombination graph (ARG) inferred from sequence data (20, 27). Here, we bypass ARG inference, which is computationally intensive and can introduce an additional layer of misspecification, by working directly with haplotype matrices. In other recent work, site frequency spectra were used as inputs to the DANN, though this summary statistic removes any linkage signal between SNPs (83). By working with haplotypes (18), we can fully leverage all available data without the need for additional inferences, making it more straightforward and better suited for hard vs soft sweep inference in aDNA. This framework is generalizable and in future work could be extended to other organisms with low coverage data or where the demography is not fully characterized.

Our findings highlight the power of deep learning for uncovering signatures of selection in aDNA data. By applying a DANN to aDNA data, we find that hard sweeps have played an important role in the evolutionary history of humans. Moreover, our approach opens the door to a range of future applications of deep learning in aDNA to study adaptation, including incorporating time as an explicit variable to examine how selection has fluctuated across historical periods or extending our approach to detect other modes of selection. In future work, ancient DNA could be used to pinpoint the specific mutations underlying putative sweeps and to narrow the set of variants prioritized for functional validation. For example, if a sweep is inferred to initiate in a particular time interval, one could focus on mutations that arose during that period or on environmental changes that plausibly altered selection pressures. As ancient DNA datasets expand and deep learning approaches for population genetics mature, it should be possible to link inferred sweeps more directly to their molecular targets and to the ecological and historical contexts in which they arose.

## 4. Methods

Detailed descriptions of data accessibility, selective sweep simulations, data processing, DANN architecture and training, and the implementation of genome-wide scans in aDNA are provided in the Supplementary Information.

## Data availability

Ancient DNA data were obtained from the Allen Ancient DNA Resource (AADR, version 51; https://dataverse.harvard.edu/dataset.xhtml?persistentId=doi:10.7910/DVN/FFIDCW). Modern human DNA data from 99 European samples (CEU) were obtained from the 1000 Genomes Project (50). Drosophila melanogaster data were downloaded from the publicly available Drosophila Genome Nexus dataset (84), which includes 205 Drosophila Genetic Reference Panel (DGRP) strains from Raleigh, North Carolina. These data can be downloaded at www.johnpool.net.

## Code availability

All computer code associated with this manuscript is available at https://github.com/garudlab/DANNsweeps.

## Acknowledgements

The authors thank Vagheesh Narasimhan and Devansh Pandey for providing processed aDNA data and for helpful conversations. The authors also thank members of the Garud Lab, particularly Peter Laurin, for helpful conversations and feedback on the paper. This work was funded by an NIGMS NIH award (R35GM151023) to NRG, an NSF CAREER award (no. 2240098) to NRG, NIGMS NIH award (R35GM127070) to AS, a UCLA Dissertation Year Award to MH, and the CSHL School of Biological Sciences Gladys & Roland Harriman Fellowship ZM.

## References

1. A. E. Page, et al., Reproductive trade-offs in extant hunter-gatherers suggest adaptive mechanism for the Neolithic expansion. Proc. Natl. Acad. Sci. U.S.A. 113, 4694–4699 (2016).

2. S. Marciniak, et al., An integrative skeletal and paleogenomic analysis of stature variation suggests relatively reduced health for early European farmers. Proc. Natl. Acad. Sci. U.S.A. 119, e2106743119 (2022).

3. Y. Souilmi, et al., Admixture has obscured signals of historical hard sweeps in humans. Nat Ecol Evol 6, 2003–2015 (2022).

4. A. Akbari, et al., Pervasive findings of directional selection realize the promise of ancient DNA to elucidate human adaptation. [Preprint] (2024). Available at: http://biorxiv.org/lookup/doi/10.1101/2024.09.14.613021 [Accessed 28 March 2025].

5. G. Kerner, et al., Genetic adaptation to pathogens and increased risk of inflammatory disorders in post-Neolithic Europe. Cell Genomics 3, 100248 (2023).

6. J. Hermisson, P. S. Pennings, Soft Sweeps: molecular population genetics of adaptation from standing genetic variation. Genetics 169, 2335–2352 (2005).

7. P. S. Pennings, J. Hermisson, Soft Sweeps II—Molecular Population Genetics of Adaptation from Recurrent Mutation or Migration. Molecular Biology and Evolution 23, 1076–1084 (2006).

8. P. W. Messer, D. A. Petrov, Population genomics of rapid adaptation by soft selective sweeps. Trends in Ecology & Evolution 28, 659–669 (2013).

9. J. Hermisson, P. S. Pennings, Soft sweeps and beyond: understanding the patterns and probabilities of selection footprints under rapid adaptation. Methods Ecol Evol 8, 700–716 (2017).

10. Y. Zheng, T. Wiehe, Adaptation in structured populations and fuzzy boundaries between hard and soft sweeps. PLOS Computational Biology 15, e1007426 (2019).

11. R. B. Harris, A. Sackman, J. D. Jensen, On the unfounded enthusiasm for soft selective sweeps II: Examining recent evidence from humans, flies, and viruses. PLOS Genetics 14, e1007859 (2018).

12. N. R. Garud, P. W. Messer, D. A. Petrov, Detection of hard and soft selective sweeps from Drosophila melanogaster population genomic data. PLOS Genetics 17, e1009373 (2021).

13. J. L. Kelley, J. Madeoy, J. C. Calhoun, W. Swanson, J. M. Akey, Genomic signatures of positive selection in humans and the limits of outlier approaches. Genome Res. 16, 980–989 (2006).

14. S. Sheehan, Y. S. Song, Deep Learning for Population Genetic Inference. PLOS Computational Biology 12, e1004845 (2016).

15. Z. Wang, et al., Automatic inference of demographic parameters using generative adversarial networks. Molecular Ecology Resources 21, 2689–2705 (2021).

16. L. Flagel, Y. Brandvain, D. R. Schrider, The Unreasonable Effectiveness of Convolutional Neural Networks in Population Genetic Inference. Molecular Biology and Evolution 36, 220–238 (2019).

17. J. R. Adrion, J. G. Galloway, A. D. Kern, Predicting the Landscape of Recombination Using Deep Learning. Molecular Biology and Evolution 37, 1790–1808 (2020).

18. L. Torada, et al., ImaGene: a convolutional neural network to quantify natural selection from genomic data. BMC Bioinformatics 20, 337 (2019).

19. A. T. Xue, et al., Discovery of Ongoing Selective Sweeps within *Anopheles* Mosquito Populations Using Deep Learning. Molecular Biology and Evolution 38, 1168–1183 (2021).

20. H. A. Hejase, Z. Mo, L. Campagna, A. Siepel, A Deep-Learning Approach for Inference of Selective Sweeps from the Ancestral Recombination Graph. Molecular Biology and Evolution 39, msab332 (2022).

21. M. R. Amin, M. Hasan, M. DeGiorgio, Digital Image Processing to Detect Adaptive Evolution. Molecular Biology and Evolution 41, msae242 (2024).

22. L. S. Whitehouse, D. R. Schrider, Timesweeper: accurately identifying selective sweeps using population genomic time series. Genetics 224, iyad084 (2023).

23. A. Dabi, D. R. Schrider, Population size rescaling significantly biases outcomes of forward-in-time population genetic simulations. Genetics 229, iyae180 (2025).

24. H. Shimodaira, Improving predictive inference under covariate shift by weighting the log-likelihood function. Journal of Statistical Planning and Inference 90, 227–244 (2000).

25. K. E. Burger, P. Pfaffelhuber, F. Baumdicker, Neural networks for self-adjusting mutation rate estimation when the recombination rate is unknown. PLOS Computational Biology 18, e1010407 (2022).

26. R. Riley, I. Mathieson, S. Mathieson, Interpreting generative adversarial networks to infer natural selection from genetic data. Genetics 226, iyae024 (2024).

27. Z. Mo, A. Siepel, Domain-adaptive neural networks improve supervised machine learning based on simulated population genetic data. PLoS Genet 19, e1011032 (2023).

28. Y. Ganin, V. Lempitsky, Unsupervised Domain Adaptation by Backpropagation. [Preprint] (2014). Available at: http://arxiv.org/abs/1409.7495 [Accessed 22 April 2025].

29. G. Csurka, “A Comprehensive Survey on Domain Adaptation for Visual Applications” in Domain Adaptation in Computer Vision Applications, G. Csurka, Ed. (Springer International Publishing, 2017), pp. 1–35.

30. V. M. Patel, R. Gopalan, R. Li, R. Chellappa, Visual Domain Adaptation: A survey of recent advances. IEEE Signal Processing Magazine 32, 53–69 (2015).

31. F. Nourbakhsh, E. Granger, G. Fumera, An Extended Sparse Classification Framework for Domain Adaptation in Video Surveillance in Computer Vision – ACCV 2016 Workshops, C.-S. Chen, J. Lu, K.-K. Ma, Eds. (Springer International Publishing, 2017), pp. 360–376.

32. A. Ramponi, B. Plank, Neural Unsupervised Domain Adaptation in NLP—A Survey in Proceedings of the 28th International Conference on Computational Linguistics, (International Committee on Computational Linguistics, 2020), pp. 6838–6855.

33. M. Rostami, D. Bose, S. Narayanan, A. Galstyan, Domain Adaptation for Sentiment Analysis Using Robust Internal Representations in Findings of the Association for Computational Linguistics: EMNLP 2023, (Association for Computational Linguistics, 2023), pp. 11484–11498.

34. K. Cochran, et al., Domain-adaptive neural networks improve cross-species prediction of transcription factor binding. Genome Res. 32, 512–523 (2022).

35. I. Mathieson, et al., Genome-wide patterns of selection in 230 ancient Eurasians. Nature 528, 499–503 (2015).

36. S. Brace, et al., Ancient genomes indicate population replacement in Early Neolithic Britain. Nature Ecology & Evolution 3, 765–771 (2019).

37. D. M. Fernandes, et al., The spread of steppe and Iranian-related ancestry in the islands of the western Mediterranean. Nature Ecology & Evolution 4, 334–345 (2020).

38. É. Harney, et al., A minimally destructive protocol for DNA extraction from ancient teeth. Genome Res. 31, 472–483 (2021).

39. M. Lipson, et al., Parallel palaeogenomic transects reveal complex genetic history of early European farmers. Nature 551, 368–372 (2017).

40. V. M. Narasimhan, et al., The formation of human populations in South and Central Asia. Science 365, eaat7487 (2019).

41. I. Olalde, et al., The genomic history of the Iberian Peninsula over the past 8000 years. Science 363, 1230–1234 (2019).

42. I. Olalde, et al., The Beaker phenomenon and the genomic transformation of northwest Europe. Nature 555, 190–196 (2018).

43. L. Papac, et al., Dynamic changes in genomic and social structures in third millennium BCE central Europe. Science Advances 7, eabi6941.

44. N. Patterson, et al., Large-scale migration into Britain during the Middle to Late Bronze Age. Nature 601, 588–594 (2022).

45. N. O’Sullivan, et al., Ancient genome-wide analyses infer kinship structure in an Early Medieval Alemannic graveyard. Science Advances 4, eaao1262.

46. V. Villalba-Mouco, et al., Survival of Late Pleistocene Hunter-Gatherer Ancestry in the Iberian Peninsula. Current Biology 29, 1169–1177.e7 (2019).

47. M. Novak, et al., Genome-wide analysis of nearly all the victims of a 6200 year old massacre. PLOS ONE 16, e0247332 (2021).

48. D. Pandey, M. Harris, N. R. Garud, V. M. Narasimhan, Leveraging ancient DNA to uncover signals of natural selection in Europe lost due to admixture or drift. Nat Commun 15, 9772 (2024).

49. P. Skoglund, I. Mathieson, Ancient Genomics of Modern Humans: The First Decade. Annu. Rev. Genom. Hum. Genet. 19, 381–404 (2018).

50. The 1000 Genomes Project Consortium, et al., A global reference for human genetic variation. Nature 526, 68–74 (2015).

51. H. Zhao, N. Alachiotis, Data preprocessing methods for selective sweep detection using convolutional neural networks. Methods 233, 19–29 (2025).

52. R. M. Cecil, L. A. Sugden, On convolutional neural networks for selection inference: Revealing the effect of preprocessing on model learning and the capacity to discover novel patterns. PLoS Comput Biol 19, e1010979 (2023).

53. P. D. B. Damgaard, et al., 137 ancient human genomes from across the Eurasian steppes. Nature 557, 369–374 (2018).

54. E. R. Jones, et al., Upper Palaeolithic genomes reveal deep roots of modern Eurasians. Nat Commun 6, 8912 (2015).

55. J. Kamm, J. Terhorst, R. Durbin, Y. S. Song, Efficiently inferring the demographic history of many populations with allele count data. J. Am. Stat. Assoc. 115 (2019).

56. N. R. Garud, P. W. Messer, E. O. Buzbas, D. A. Petrov, Recent Selective Sweeps in North American Drosophila melanogaster Show Signatures of Soft Sweeps. PLoS Genet 11, e1005004 (2015).

57. E. K. Irving-Pease, et al., The selection landscape and genetic legacy of ancient Eurasians. Nature 625, 312–320 (2024).

58. M. K. Le, et al., 1,000 ancient genomes uncover 10,000 years of natural selection in Europe. BioRxiv (2022). 10.1101/2022.08.24.505188.

59. E. M. Leffler, et al., Multiple Instances of Ancient Balancing Selection Shared Between Humans and Chimpanzees. Science 339, 1578–1582 (2013).

60. P. I. W. de Bakker, S. Raychaudhuri, Interrogating the major histocompatibility complex with high-throughput genomics. Human Molecular Genetics 21, R29–R36 (2012).

61. T. Bersaglieri, et al., Genetic Signatures of Strong Recent Positive Selection at the Lactase Gene. The American Journal of Human Genetics 74, 1111–1120 (2004).

62. N. S. Enattah, et al., Identification of a variant associated with adult-type hypolactasia. Nature Genetics 30, 233–237 (2002).

63. I. Mathieson, J. Terhorst, Direct detection of natural selection in Bronze Age Britain. Genome Res. 32, 2057–2067 (2022).

64. L. Segurel, et al., Why and when was lactase persistence selected for? Insights from Central Asian herders and ancient DNA. PLOS Biology 18, e3000742 (2020).

65. R. P. Evershed, et al., Dairying, diseases and the evolution of lactase persistence in Europe. Nature 608, 336–345 (2022).

66. K. Watanabe, E. Taskesen, A. van Bochoven, D. Posthuma, Functional mapping and annotation of genetic associations with FUMA. Nature Communications 8, 1826 (2017).

67. D. R. Schrider, F. K. Mendes, M. W. Hahn, A. D. Kern, Soft Shoulders Ahead: Spurious Signatures of Soft and Partial Selective Sweeps Result from Linked Hard Sweeps. Genetics 200, 267–284 (2015).

68. R. E. Green, et al., A Draft Sequence of the Neandertal Genome. Science 328, 710–722 (2010).

69. C. Evers, et al., Mosaic deletion of EXOC6B: further evidence for an important role of the exocyst complex in the pathogenesis of intellectual disability. Am J Med Genet A 164A, 3088–94 (2014).

70. R. D. Hernandez, et al., Classic Selective Sweeps Were Rare in Recent Human Evolution. Science 331, 920–924 (2011).

71. D. R. Schrider, A. D. Kern, Soft Sweeps Are the Dominant Mode of Adaptation in the Human Genome. Molecular Biology and Evolution 34, 1863–1877 (2017).

72. M. DeGiorgio, C. D. Huber, M. J. Hubisz, I. Hellmann, R. Nielsen, SweepFinder2: increased sensitivity, robustness and flexibility. Bioinformatics 32, 1895–1897 (2016).

73. P. S. Pennings, J. Hermisson, Soft Sweeps III: The Signature of Positive Selection from Recurrent Mutation. PLOS Genetics 2, e186 (2006).

74. T. Karasov, P. W. Messer, D. A. Petrov, Evidence that Adaptation in Drosophila Is Not Limited by Mutation at Single Sites. PLOS Genetics 6, e1000924 (2010).

75. A. Mutero, M. Pralavorio, J. M. Bride, D. Fournier, Resistance-associated point mutations in insecticide-insensitive acetylcholinesterase. Proceedings of the National Academy of Sciences 91, 5922–5926 (1994).

76. P. Menozzi, M. A. Shi, A. Lougarre, Z. H. Tang, D. Fournier, Mutations of acetylcholinesterase which confer insecticide resistance in Drosophila melanogaster populations. BMC Evolutionary Biology 4, 4 (2004).

77. M. M. Magwire, F. Bayer, C. L. Webster, C. Cao, F. M. Jiggins, Successive Increases in the Resistance of Drosophila to Viral Infection through a Transposon Insertion Followed by a Duplication. PLOS Genetics 7, e1002337 (2011).

78. P. Daborn, S. Boundy, J. Yen, B. Pittendrigh, R. ffrench-Constant, DDT resistance in Drosophila correlates with Cyp6g1 over-expression and confers cross-resistance to the neonicotinoid imidacloprid. Molecular Genetics and Genomics 266, 556–563 (2001).

79. J. M. Schmidt, et al., Copy Number Variation and Transposable Elements Feature in Recent, Ongoing Adaptation at the Cyp6g1 Locus. PLOS Genetics 6, e1000998 (2010).

80. Y. T. Aminetzach, J. M. Macpherson, D. A. Petrov, Pesticide Resistance via Transposition-Mediated Adaptive Gene Truncation in Drosophila. Science 309, 764–767 (2005).

81. S. Mallick, D. Reich, The Allen Ancient DNA Resource (AADR): A curated compendium of ancient human genomes. Harvard Dataverse. 10.7910/DVN/FFIDCW. Deposited 2023.

82. S. Mallick, et al., The Allen Ancient DNA Resource (AADR) a curated compendium of ancient human genomes. Scientific Data 11, 182 (2024).

83. K. A. Cobb, M. L. Smith, The reasonable effectiveness of domain adaptation for inference of introgression. [Preprint] (2025). Available at: http://biorxiv.org/lookup/doi/10.1101/2025.01.17.633659 [Accessed 25 August 2025].

84. J. B. Lack, et al., The Drosophila Genome Nexus: A Population Genomic Resource of 623 Drosophila melanogaster Genomes, Including 197 from a Single Ancestral Range Population. Genetics 199, 1229–1241 (2015).

