## Supplementary Information for "The persistence and loss of hard selective sweeps amid admixture in ancient Eurasians"

Supplementary Material

Table of Contents

#### Supplemental Figures:

**Figure S1.** Performance of DANN using different haplotype sorting approaches………………. 3

**Figure S2.** Proportion of missing data in aDNA samples………………………………………4

**Figure S3.** Misspecified admixture models to test DANN versus CNN performance……….…. 4

**Figure S4.** Performance of the DANN using a one-versus-rest approach……..…………….….. 5

**Figure S5.** Confusion matrices for the DANN trained and tested on simulated data...…………. 6

**Figure S6.** Confusion matrices for the DANN trained with aDNA………………..…………….. 6

**Figure S7.** Precision recall curves for the DANN, CNN and H12……………………...……... 7

**Figure S8.** Proportion of hard and soft sweeps detected as a function of $\theta_A$ and $s$…………...…. 8

**Figure S9.** Sweep sharing across time periods…………………………………………….…... 9

**Fig S10.** Genome-wide DANN scans across ancient time periods using a constant $N_e$ model as source domain.……………………………………………………………………….….. 11

**Fig S15.** Scan of chromosome 15 for epoch IA showing “soft shoulder effect” ……………..... 16

**Fig S16.** Confusion matrix for estimating misclassification of sweeps in the aDNA DANN…….17

**Fig S17.** Haplotype visualization of 24 sweeps from Figure 4A…..……………………………. 19

**Fig S18.** Haplotype visualization of a window classified as soft in the flanking regions of the HLA sweep in period H and a window classified as neutral……………………………………….. 20

**Table S3.** Overlap between DANN-detected sweeps and those reported in previous studies…... 26

#### Supplemental Text:

**Text S1.** Ancient DNA data ………………………………………………………………..27
**Text S2:** DANN model architecture ………………………………………………………… 27
**Text S3.** Haplotype sorting ……………………………………………………………….. 28
**Text S4:** Simulations ………………………………………………………...……… 29
**Text S5:** Model training ……………………………………………………………...30
**Text S6.** DANN genome-wide scan ……………………………………………………….32
**Text S7:** ABC for hard/soft sweep classification with H12 and H2/H1 statistics …………….. 32
**Text S8:** Jaccard index and permutation test ………………………………………………... 33
**Text S9:** Application of a DANN to a North American population of *D. melanogaster* ……… 33

**References**……………………………………………………………………………… 34

#### Supplementary Figures

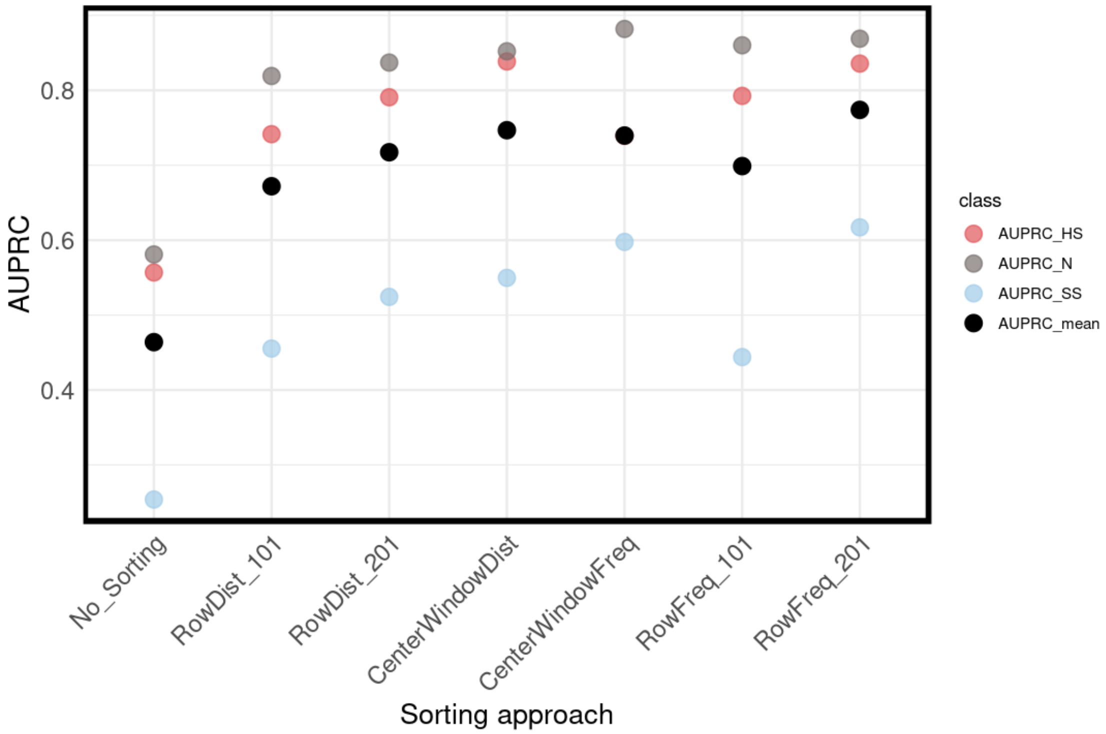

**Figure S1. Performance of DANN using different haplotype sorting approaches.** We tested three approaches for sorting haplotype images: sorting by distance to the most common haplotype (RowDist), sorting haplotypes by highest to lowest frequency (RowFreq), sorting a central 51 SNP window by RowDist or RowFreq within a 201 SNP window (CenterWindowDist and CenterWindowFreq, respectively). For RowDist and RowFreq we tested two different window sizes, a 201 SNP window and a 101 SNP window. For each model we evaluated the performance on each category, hard (red), soft (blue), and neutrality (gray), using a one-versus-rest approach and obtained the AUPRC for each class. We averaged the three resulting AUPRC giving the mean AUPRC shown as the black data points in the figure.

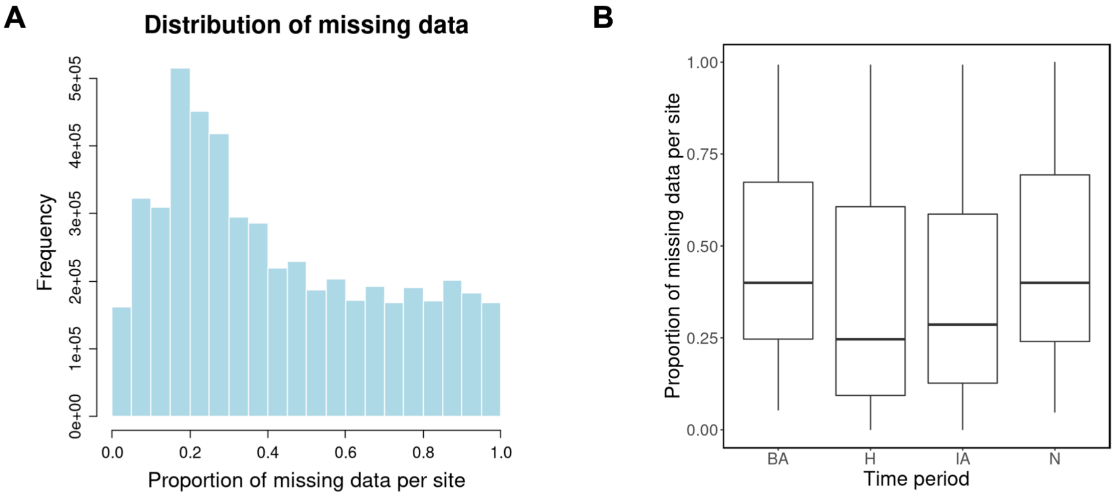

**Figure S2. Proportion of missing data in aDNA samples. (A)** Distribution of missing data across 201 SNP windows across all time periods. **(B)** Proportion of SNPs with missing data in 201 SNP windows per site for each time period. Each 201 SNP window is subsampled to 150 pseudo-haplotype with the least amount of missing data.

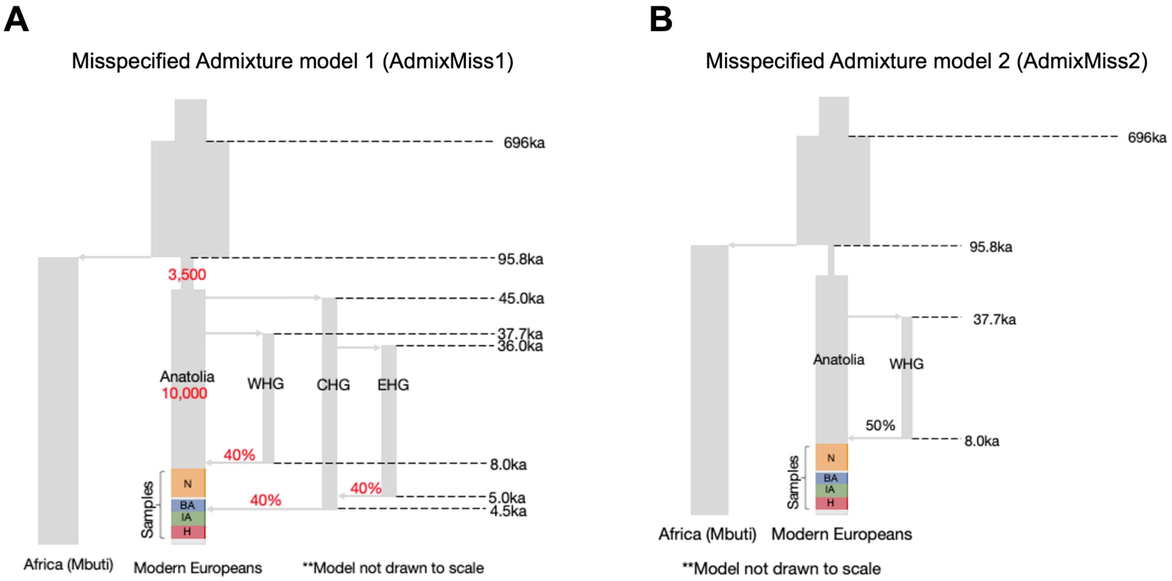

**Figure S3. Misspecified admixture models to test DANN versus CNN performance.** Figures **(A)** and **(B)** represent variations of the model shown in **Fig. 1C**. These models are used as source domains when evaluating the effectiveness of the DANN in Fig. 3. In **(A)** we highlight in red the changes to admixture proportions and the $N_e$ of the out of Africa Bottleneck and the European population branch. In **(B)** we show an admixture model in which we only include one admixing population (WHG) and remove the CHG and EHG branches.

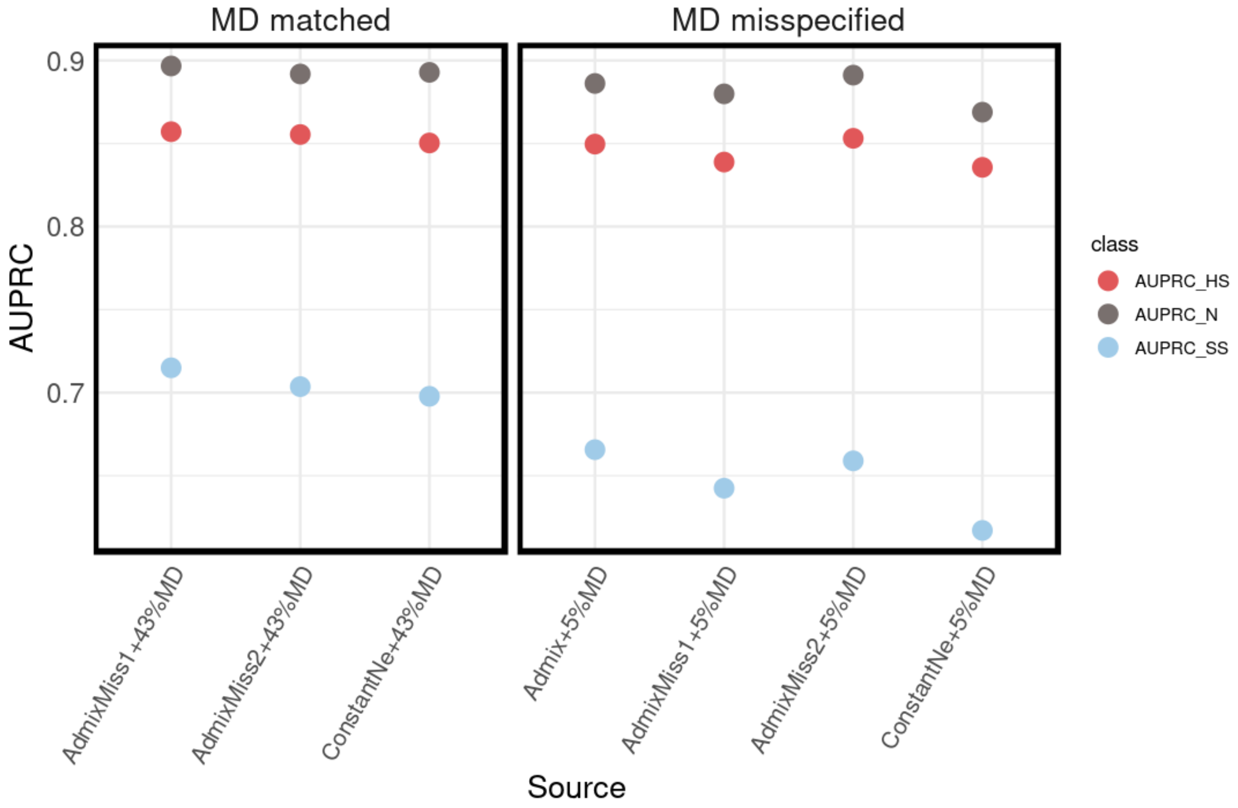

**Figure S4. Performance of the DANN using a one-versus-rest approach.** AUPRC for each category using a one-versus-rest approach. We trained a DANN setting the target domain using a human admixture model with an average of 43% missing data per site. The source domain varied in terms of demographic model used (x-axis) and proportion of missing data per site (left vs right panels). AdmixMiss1 and AdmixMiss2 refer to models drawn in Fig. S3.

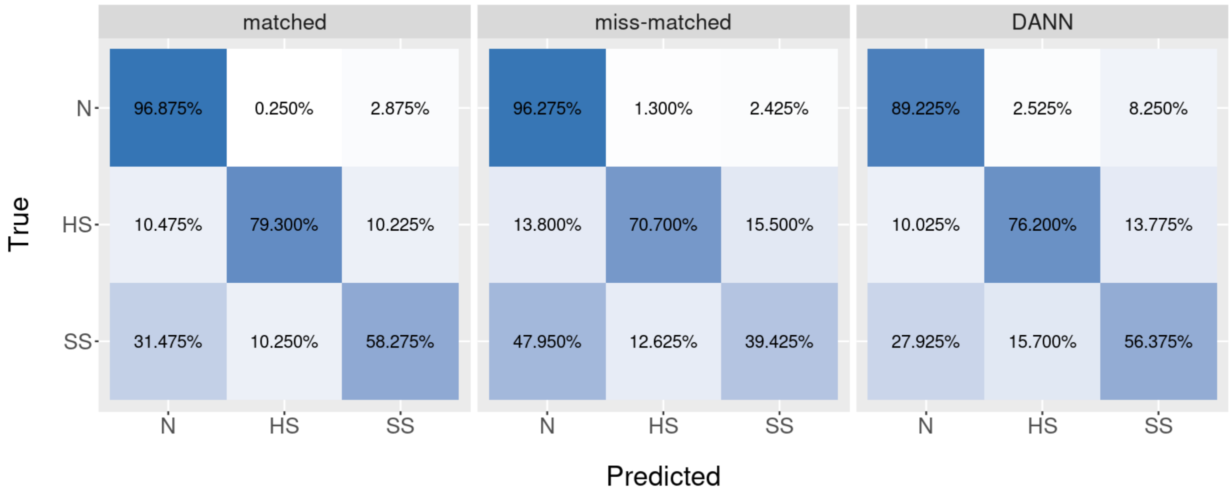

**Figure S5. Confusion matrices for the DANN trained and tested on simulated data.** (Neutral vs. hard sweep vs. soft sweep). On the left is a confusion matrix corresponding to the scenario where both source and target domains were matched and used a constant $N_e$ model with an average of 43% missing data per site. In the middle, source and target domains were mismatched where a constant $N_e$ model was used for the source domain and a human admixture model was used for the target domain, both with an average of 43% missing data per site as a target domain. On the right, the same mismatch scenario was used, but this time a DANN was turned on to correct for mismatch.

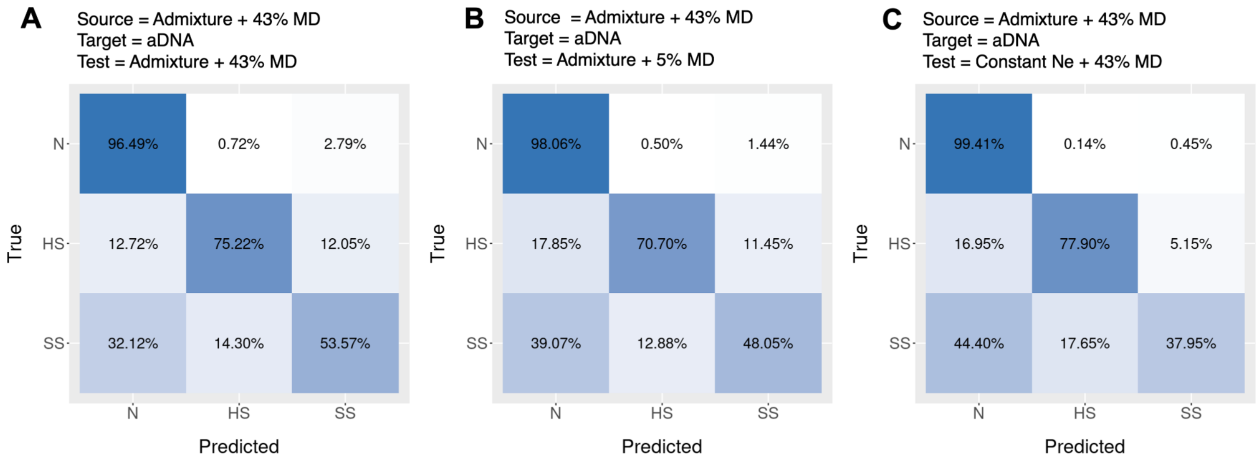

**Figure S6. Confusion matrices for the DANN trained with aDNA.** We trained the DANN on aDNA data as the target domain and the admixture model with 43% of missing data as the source domain. We tested this model in three different simulated scenarios: **(A)** Admixture model with 43% missing data (same as source domain used for training) **(B)** Admixture model with 5% missing data and **(C)** Constant $N_e$ model with 43 % missing data.

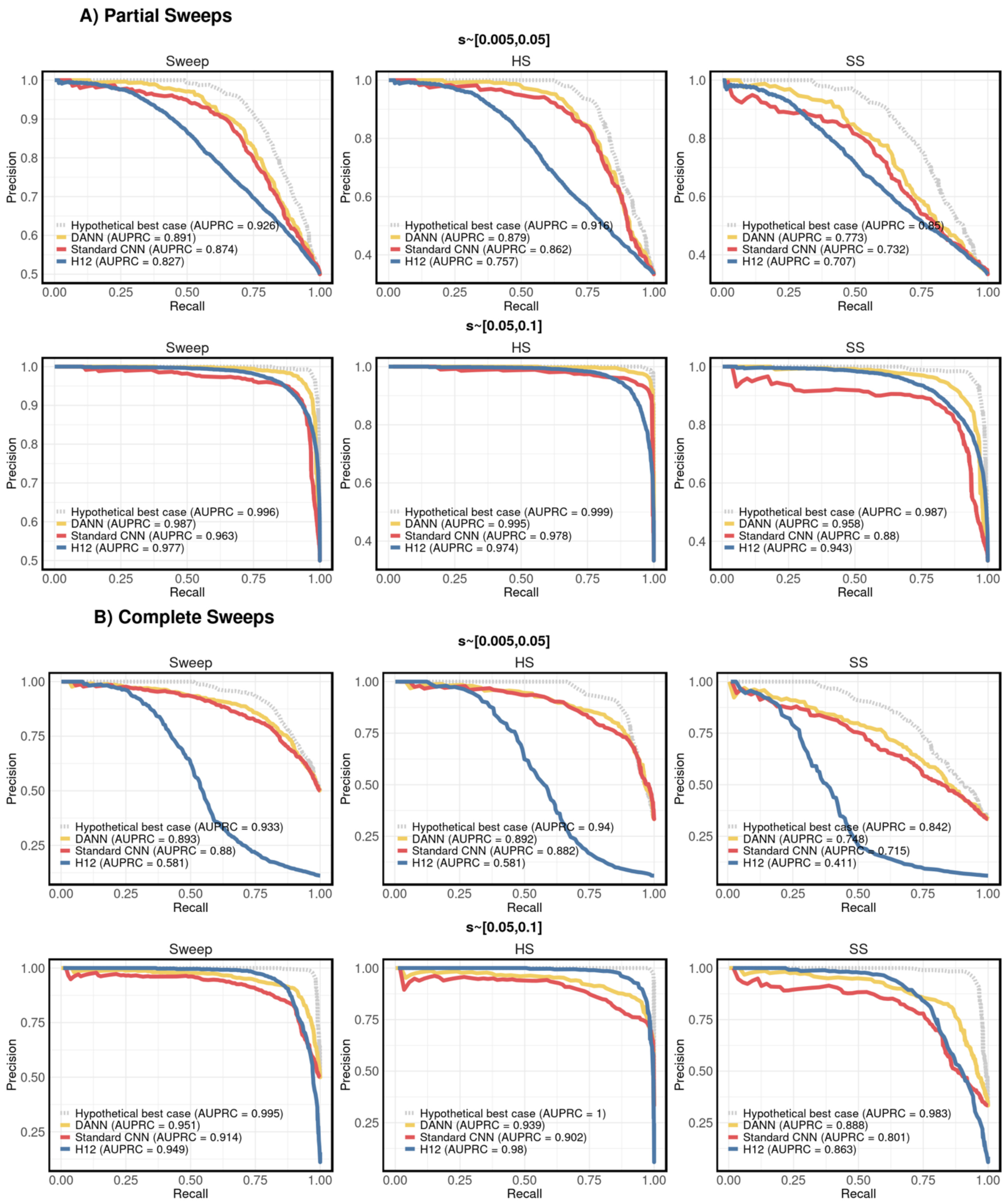

**Figure S7. Precision recall curves for the DANN, CNN and H12.** Precision recall curves for sweeps vs neutral, hard sweeps vs neutral and soft sweeps vs neutral in partial **(A)** and complete **(B)** sweeps. We test all models in a weak selection regime (s~[0.005,0.05]) and strong selection regime (s~[0.05,0.1]). The performance of the DANN is shown in yellow. The blue curve corresponds to the performance of a standard CNN tested on a matched scenario, that is tested on source domain data (i.e. constant $N_e$ model). The red curve corresponds to the performance of a standard CNN tested on a mis-specified scenario, that is tested on data from the target domain (i.e. admixture model). The grey curve shows the performance of H12 (Garud et al. 2015), a statistic that we previously applied to the same aDNA data (Harris and Pandey et al. 2024). For all scenarios evaluated, we included 500 simulations of hard sweeps, soft sweeps, and neutrality, respectively.

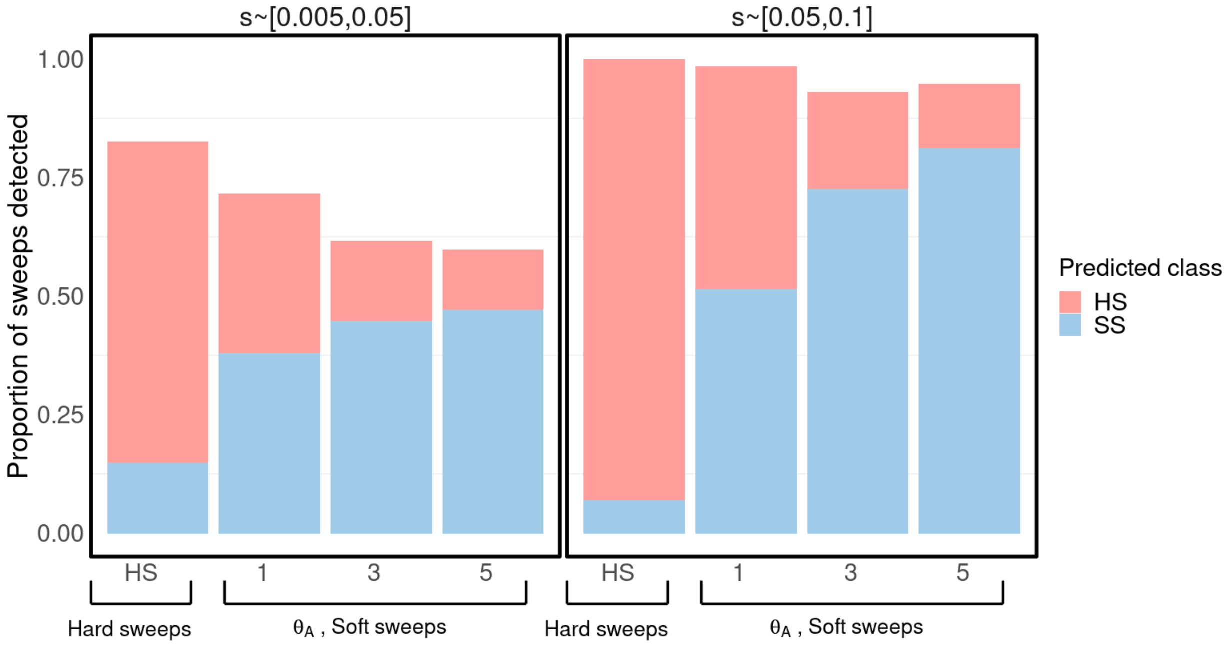

**Figure S8. Proportion of hard and soft sweeps detected as a function of $\theta_A$ and *s*.** Here the source domain was a constant *Ne* model with a 43% missing data rate and the target domain was the human admixture model with a 43% missing data rate. We test the DANN in a weak selection regime (left, s~[0.005,0.05]) and strong selection regime (right, s~[0.05,0.1]) as a function of the softness of the sweep. The y-axis shows the proportion of sweeps the DANN is able to recover, colored by the class it predicts with blue representing predicted soft sweeps and red representing predicted hard sweeps. The x-axis shows the true sweep class with hard sweeps at the left and soft sweeps at the right increasing in softness from $\theta_A = 1$ to 5.

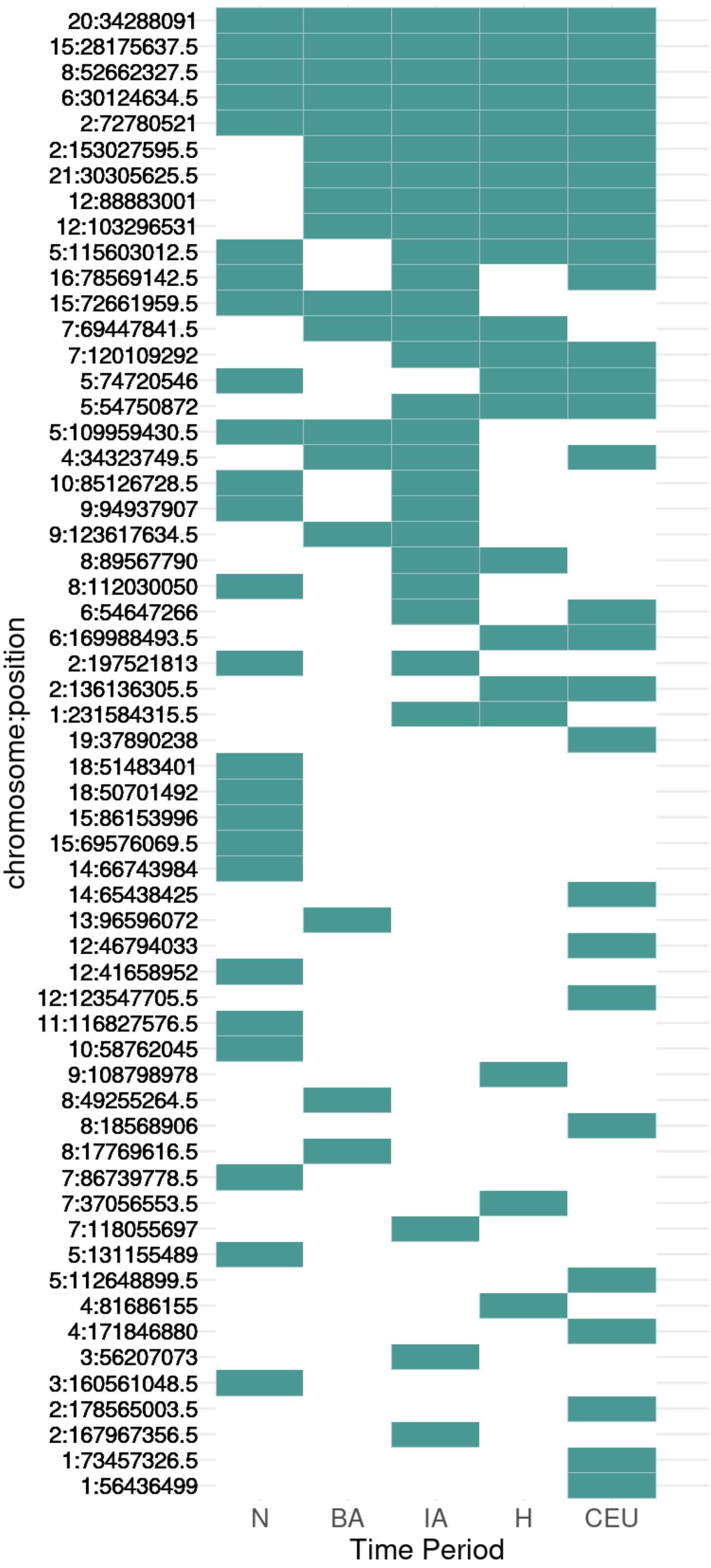

**Figure S9. Sweep sharing across time periods.** We show the occurrence of sweeps in aDNA and modern human DNA (CEU) and sort the sweeps by amount of sharing across time periods.

The coordinates of the central SNP of the analysis window with the highest probability of supporting the sweep is shown on left. A green rectangle indicates that a sweep was detected at a given time period. At top are sweeps present in all time periods and at bottom are unique sweeps found in only one time period.

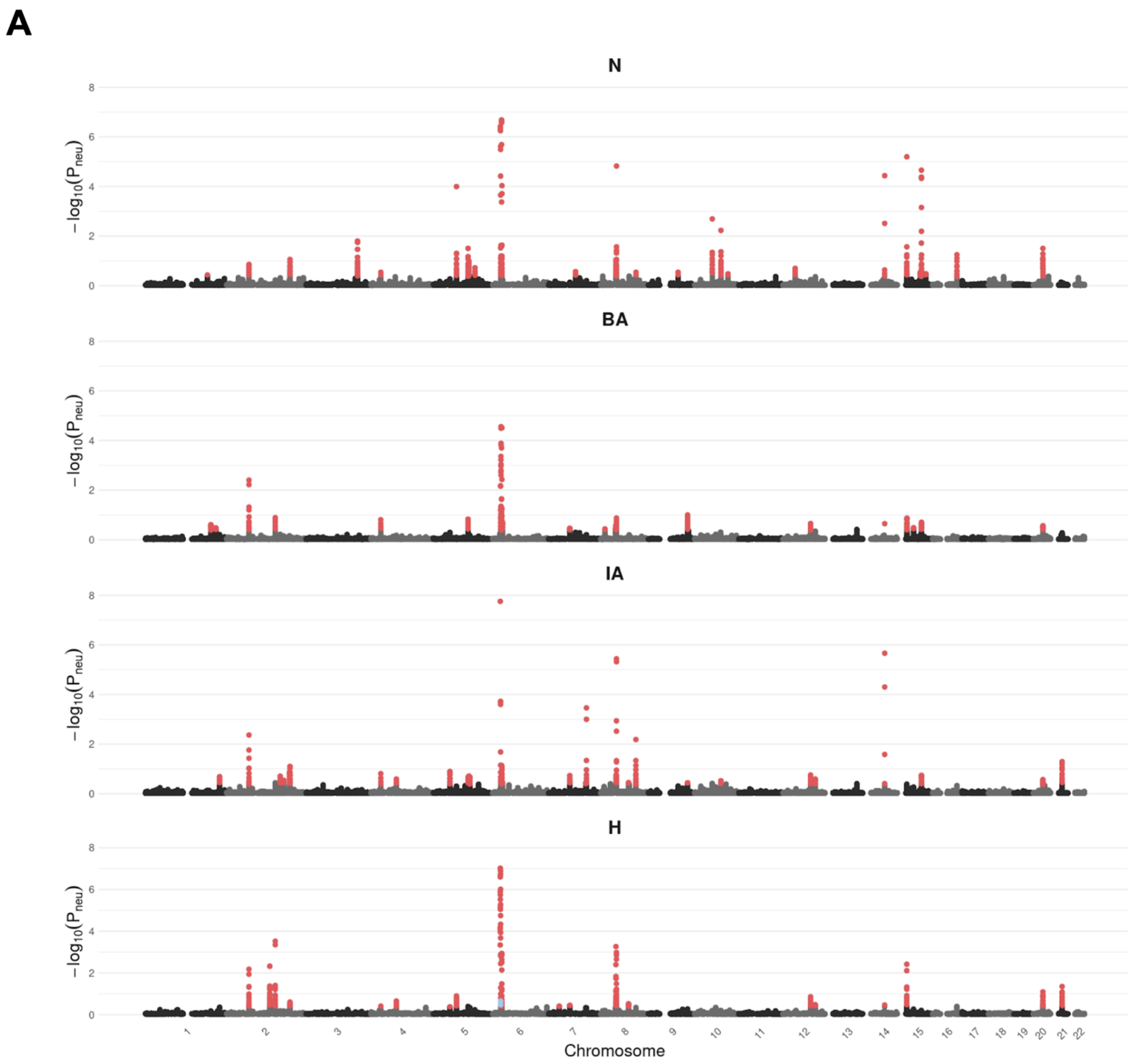

**B**

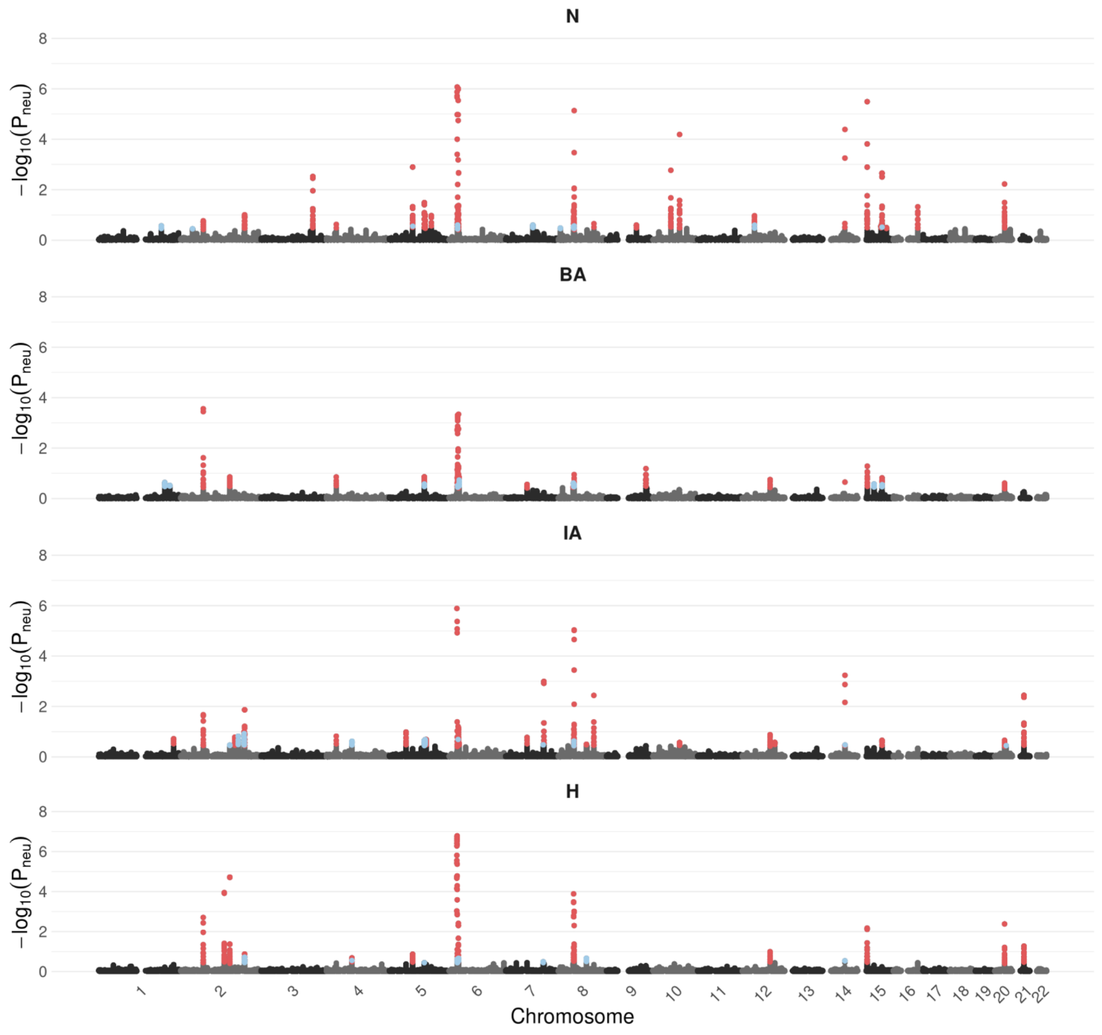

**Fig S10. Genome-wide DANN scans across ancient time periods using a constant $N_e$ model as source domain.** Unlike in Fig. 4 where the DANN was trained with an admixture model with 43% missing data as a source domain, here a constant $N_e = 10^4$ model with 43% missing data was used for the source domain. The y-axis shows the probability of neutrality -log($P_{neu}$) predicted using the DANN. Windows predicted as hard sweeps are colored in red and windows predicted as soft sweeps are colored in blue. **(A)** Soft sweep simulations were generated using a recurrent *de novo* mutation model drawing from $\Theta_A$ ~U[1,5] (see Methods). **(B)** Soft sweep simulations were generated from standing genetic variation, drawing an initial allele frequency of $PF_{init}$ ~ U(0.01,0.1) before the onset of selection.

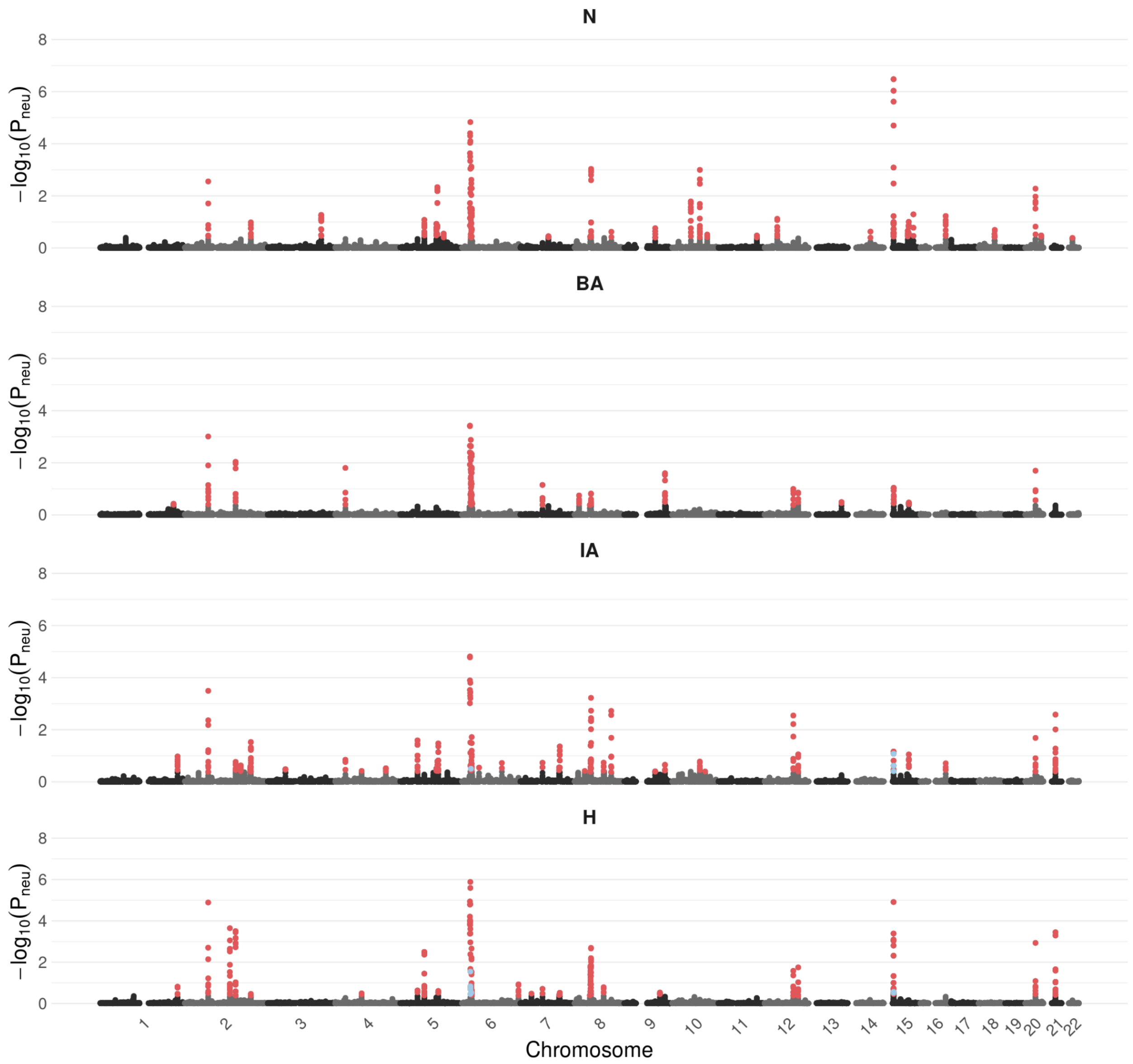

**Fig S11. Genome-wide DANN scans across ancient time periods using a human admixture model with 5% missing data as source domain.** Unlike in Figure 4 where the DANN was trained with an admixture model with 43% missing data as a source domain, here an admixture model with 5% missing data was used for the source domain and the aDNA data as target domain. The y-axis shows the probabilities of neutrality -log($P_{neu}$) predicted using the DANN. Windows predicted as hard sweeps are colored in red and windows predicted to be soft are colored in blue.

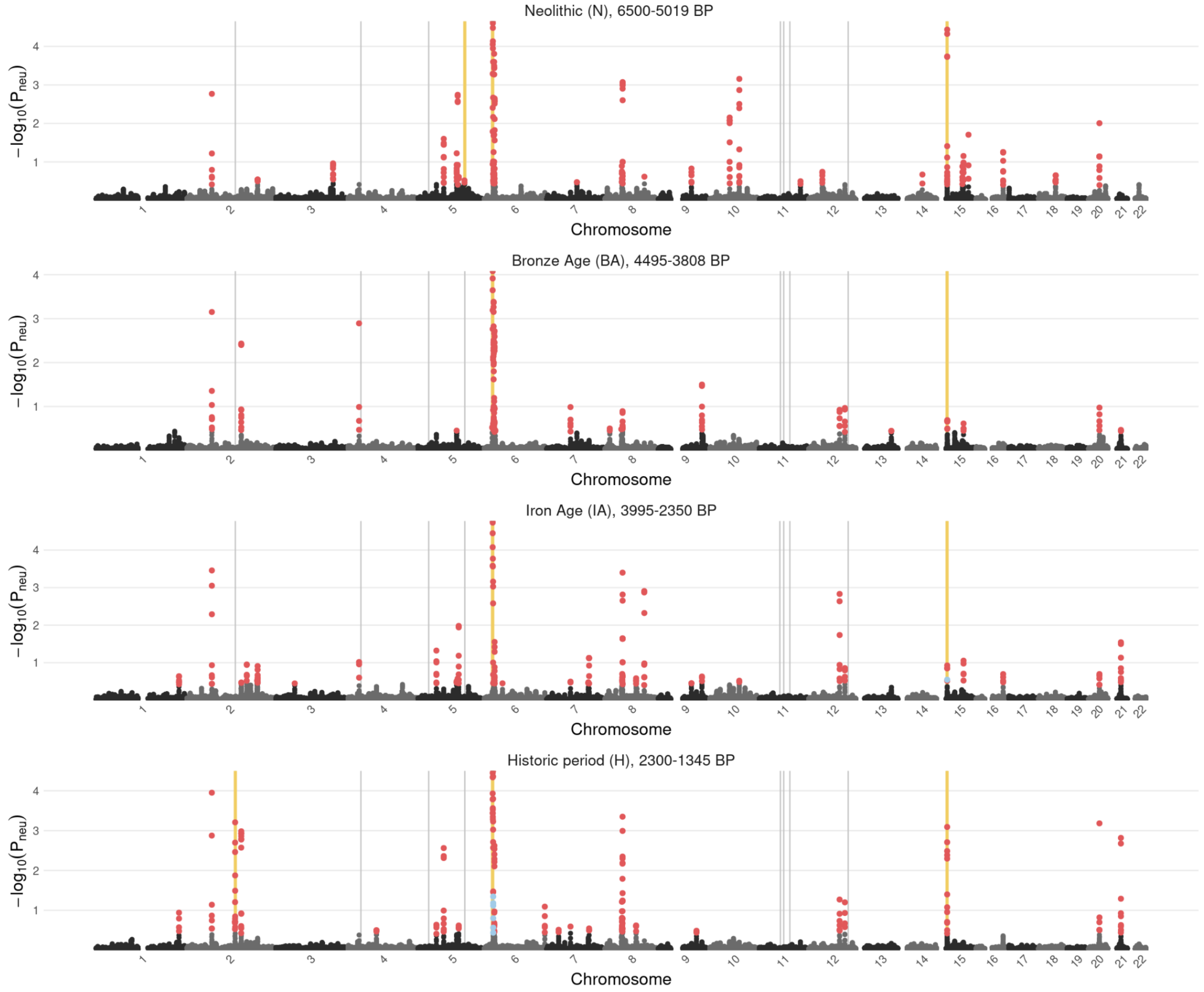

**Fig S12. Overlap with Mathieson et al. 2015.** This scan is replicated from Figure 4. The vertical lines (grey + yellow) represent the sweeps identified in Mathieson et al. 2015. Yellow vertical lines represent sweeps that overlap between our scan and the Mathieson et al. 2015 scan.

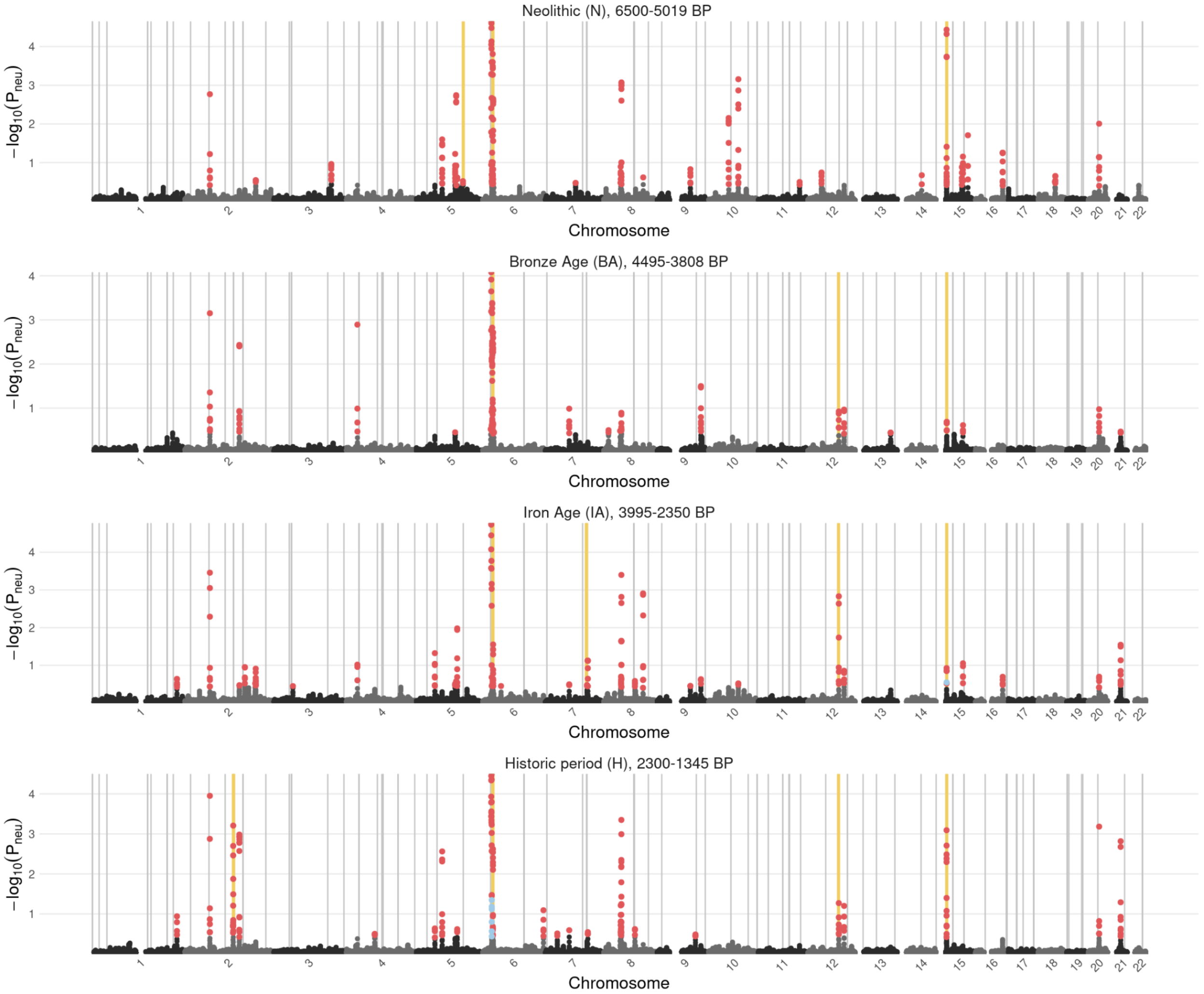

**Fig S13. Overlap with Akbari et al 2024.** This scan is replicated from Figure 4. The vertical lines (grey + yellow) represent the genes highlighted in Figure 2 of Akbari et al 2024. Yellow vertical lines represent sweeps that overlap between our scan and the Akbari et al. 2024 scan.

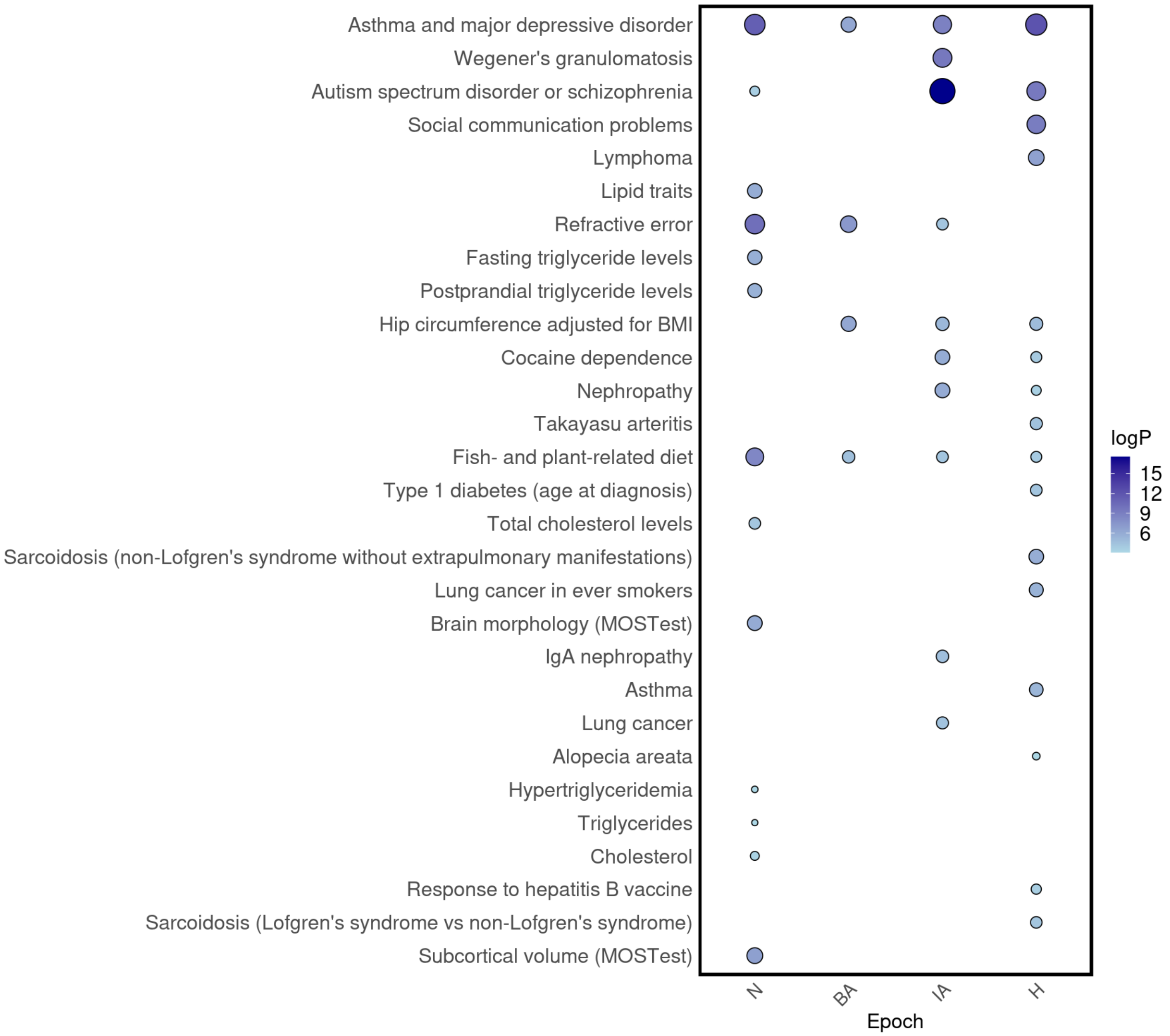

**Figure S14. Gene set enrichment analysis across epochs.** Enrichment analysis results of protein coding genes overlapping loci observed in association studies and sweep candidates detected with the DANN. Results with -logP >3 are shown above. The color and size of the circles correspond to the logP value.

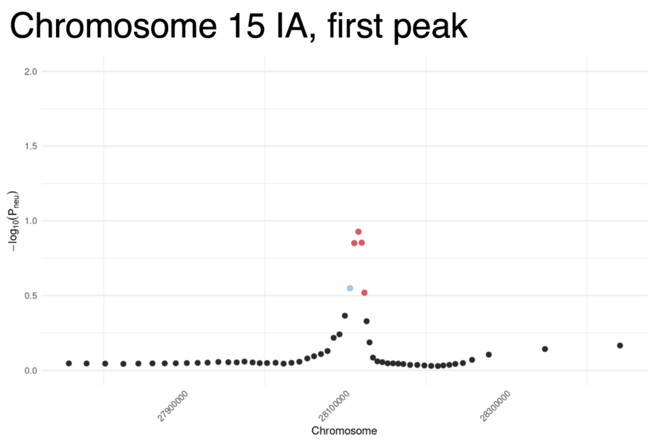

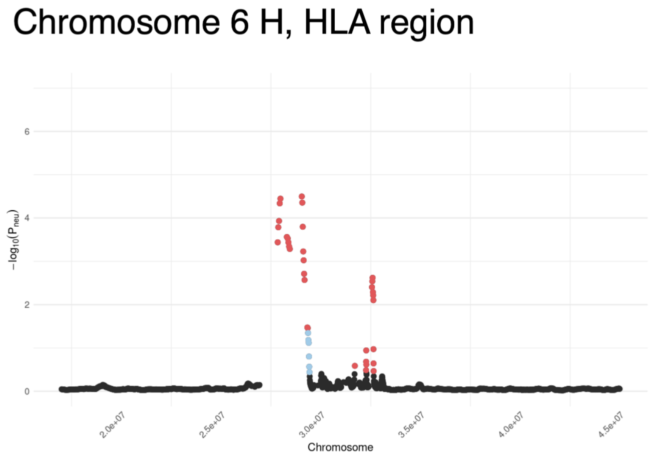

**Figure S15. Scan of chromosome 15 for epoch IA showing “soft shoulder effect”.** Colored in red are windows classified as hard sweeps and in blue are windows classified as soft sweeps. Black dots correspond to windows classified as neutral.

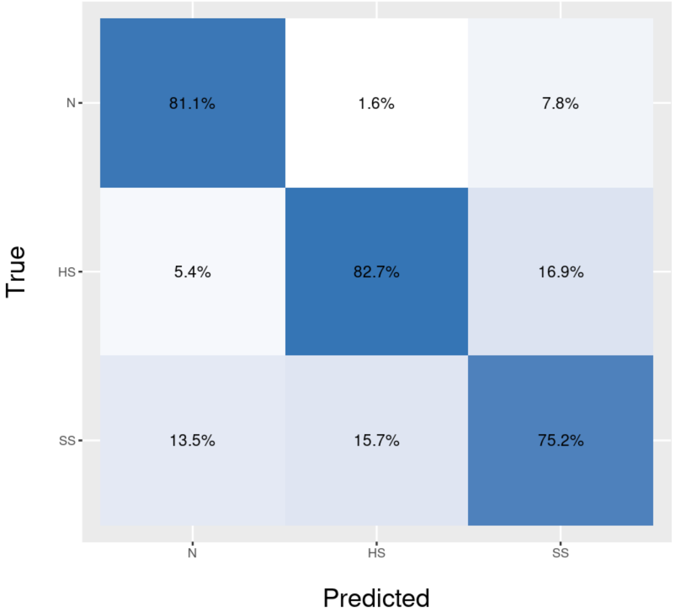

**Figure S16. Confusion matrix for estimating misclassification of sweeps in the aDNA DANN.** We test the DANN used for Fig.4A, trained on aDNA as target domain, on simulations generated under the admixture model with 43% missing data. The confusion matrix is normalized by columns to show the estimate of false discovery rates (FDR) per class.

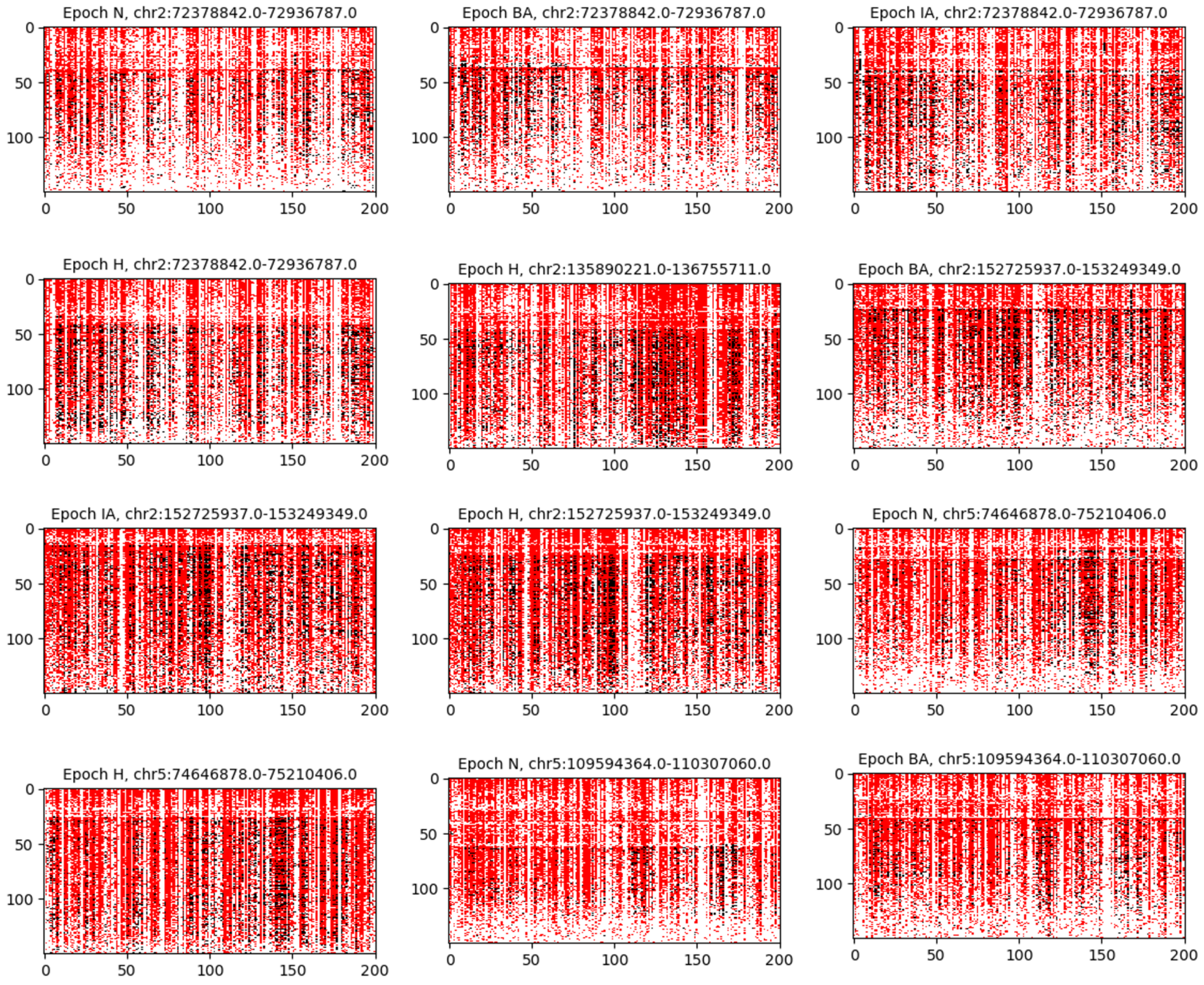
Epoch N, chr2:72378842.0-72936787.0
Epoch BA, chr2:72378842.0-72936787.0
Epoch IA, chr2:72378842.0-72936787.0
Epoch H, chr2:72378842.0-72936787.0
Epoch H, chr2:135890221.0-136755711.0
Epoch BA, chr2:152725937.0-153249349.0
Epoch IA, chr2:152725937.0-153249349.0
Epoch H, chr2:152725937.0-153249349.0
Epoch N, chr5:74646878.0-75210406.0
Epoch H, chr5:74646878.0-75210406.0
Epoch N, chr5:109594364.0-110307060.0
Epoch BA, chr5:109594364.0-110307060.0

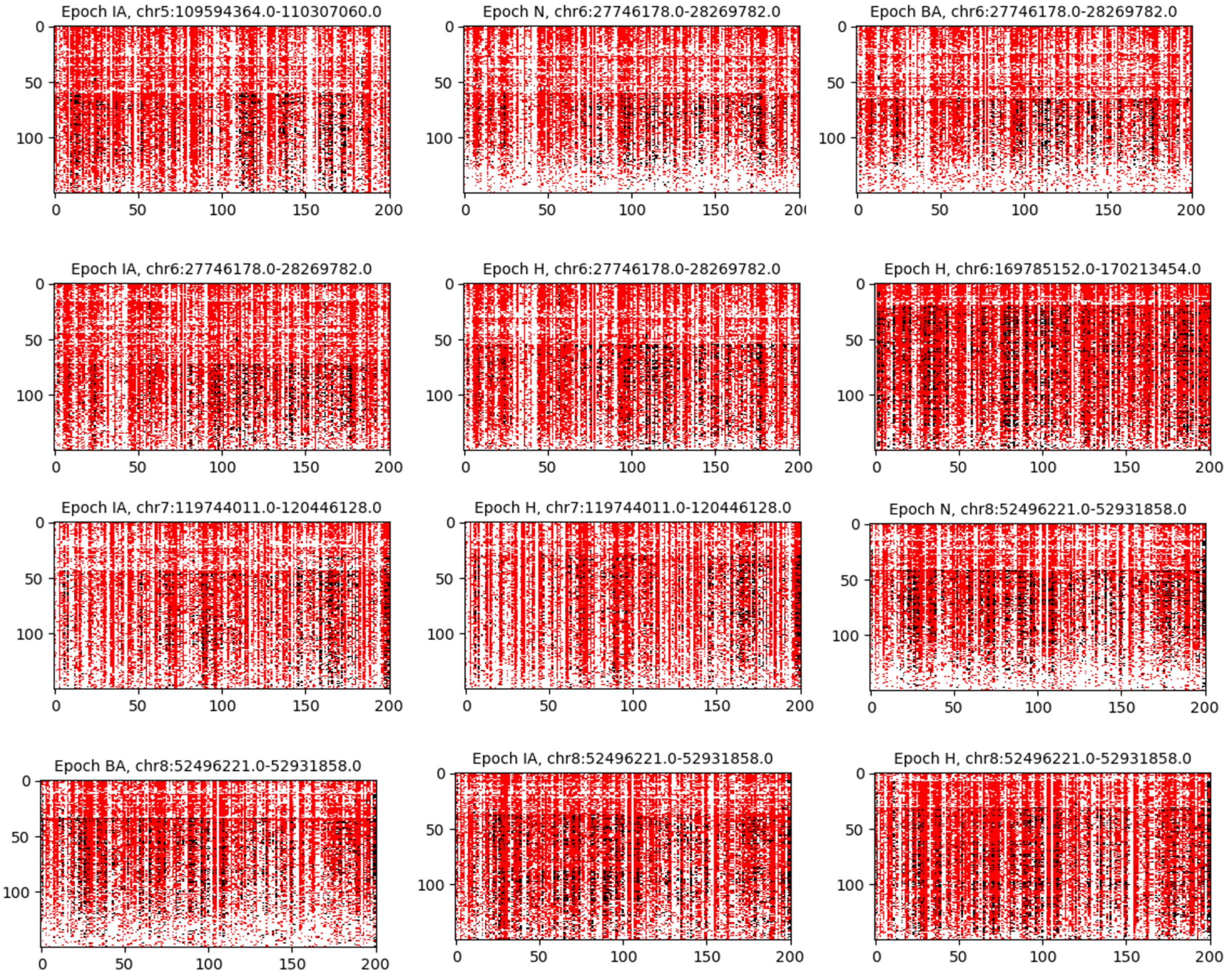

**Fig S17. Haplotype visualization of 24 sweeps from Figure 4A**. Shown are haplotype plots for the central window consisting of 201 SNPs of 24 random sweeps in Figure 4A. Haplotypes are ordered from most to least frequent. Red indicates the major allele found in the most frequent haplotype. Black indicates any difference with the allele identified in the top haplotype. White indicates missing data.

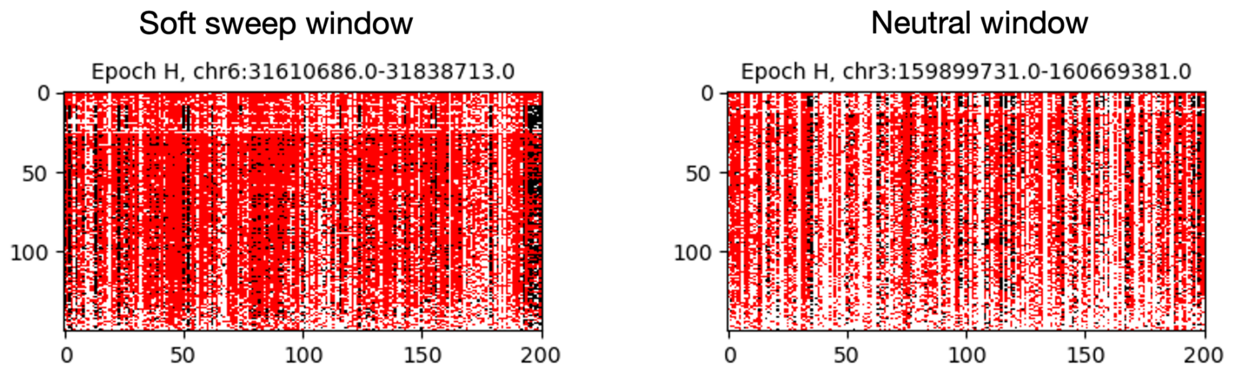

**Fig S18. Haplotype visualization of a window classified as soft in the flanking regions of the HLA sweep in period H (left) and a window classified as neutral (right).** Haplotypes are ordered from most to least frequent. Red indicates the major allele found in the most frequent haplotype. Black indicates any difference with the allele identified in the top haplotype. White indicates missing data.

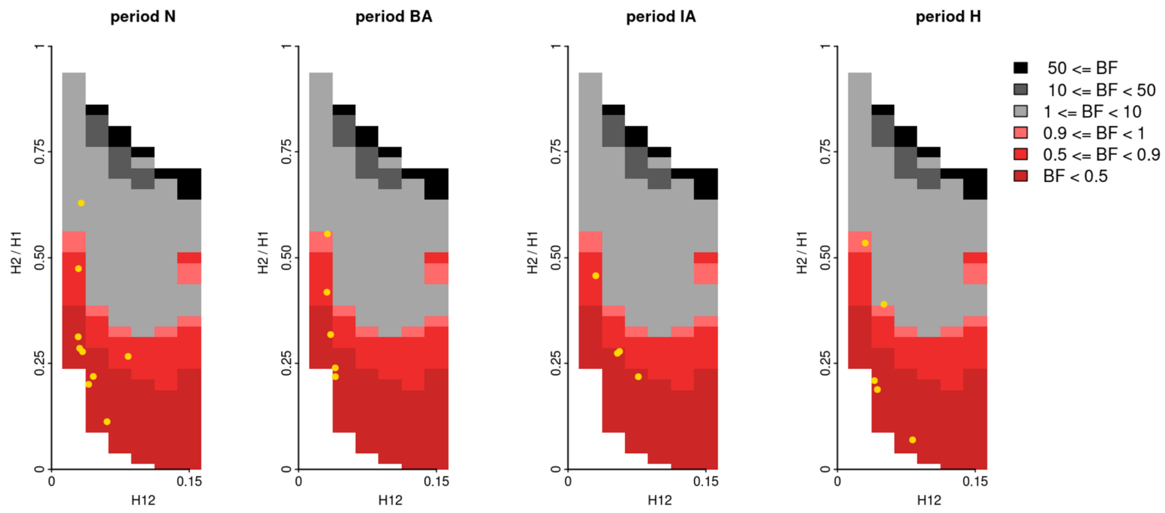

**Fig S19. Range of H12 and H2/H1 values expected to generate hard and soft sweeps from simulations under a human admixture model.** Bayes factors (BF) were calculated for a grid of H12 and H2/H1 values. In each panel we show the sweeps for each time period with H12 values greater than the 95th percentile of the values obtained from 400,000 neutral admixture simulations. In total there were 23 sweep candidates passing this 95th percentile threshold. BFs were calculated by taking the ratio of the number of soft sweep versus hard sweep simulations that were within a Euclidean distance of 10% of a given pair of H12 and H2/H1 values. Values greater than 1 indicate support for soft sweeps and values less than 1 indicate support for hard sweeps. Thus, regions colored in red represent (H12, H2/H1) pairs that are more likely to be generated from a hard sweep, while grey regions represent (H12, H2/H1) pairs that are more likely to be generated from a soft sweep. The darker the shade of the color the stronger the support for each sweep type.

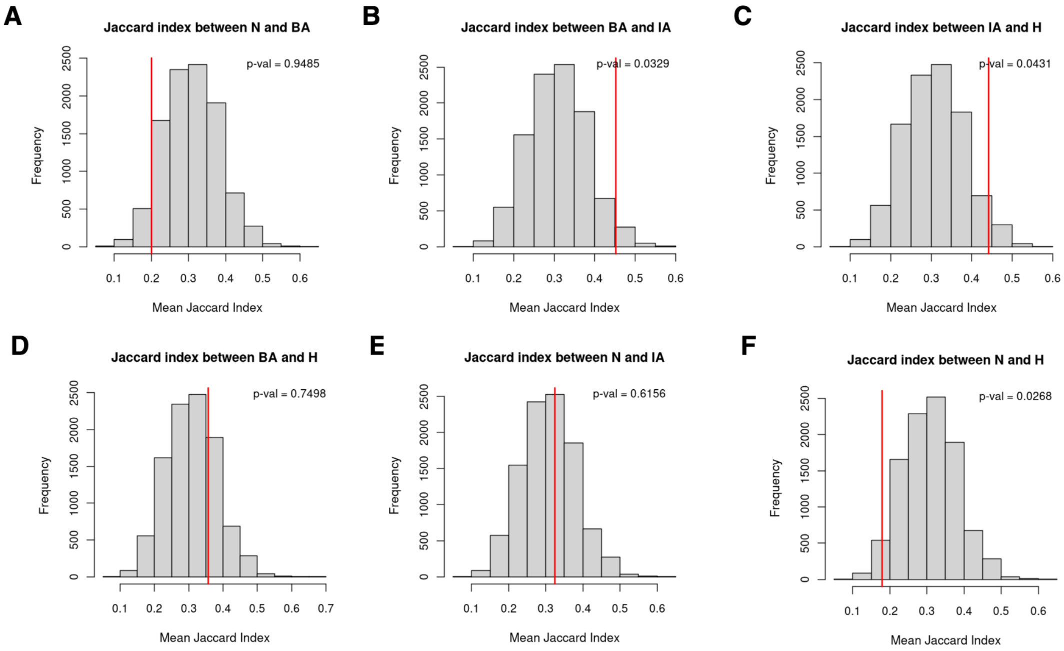

**Fig S20. Jaccard index permutation test for overlap of selective sweeps across time periods.** We show the null distributions of the Jaccard index after permuting the time period labels in a single permutation scheme. This permutation was performed 5,000 times, and Jaccard indices were measured for each of the 6 pairs of time periods for each permutation. The red vertical line shows the observed Jaccard index between two time periods. In (A-C) we test for a higher proportion of sweep sharing between time periods and in (D-F) we test for a lower proportion of sweep sharing.

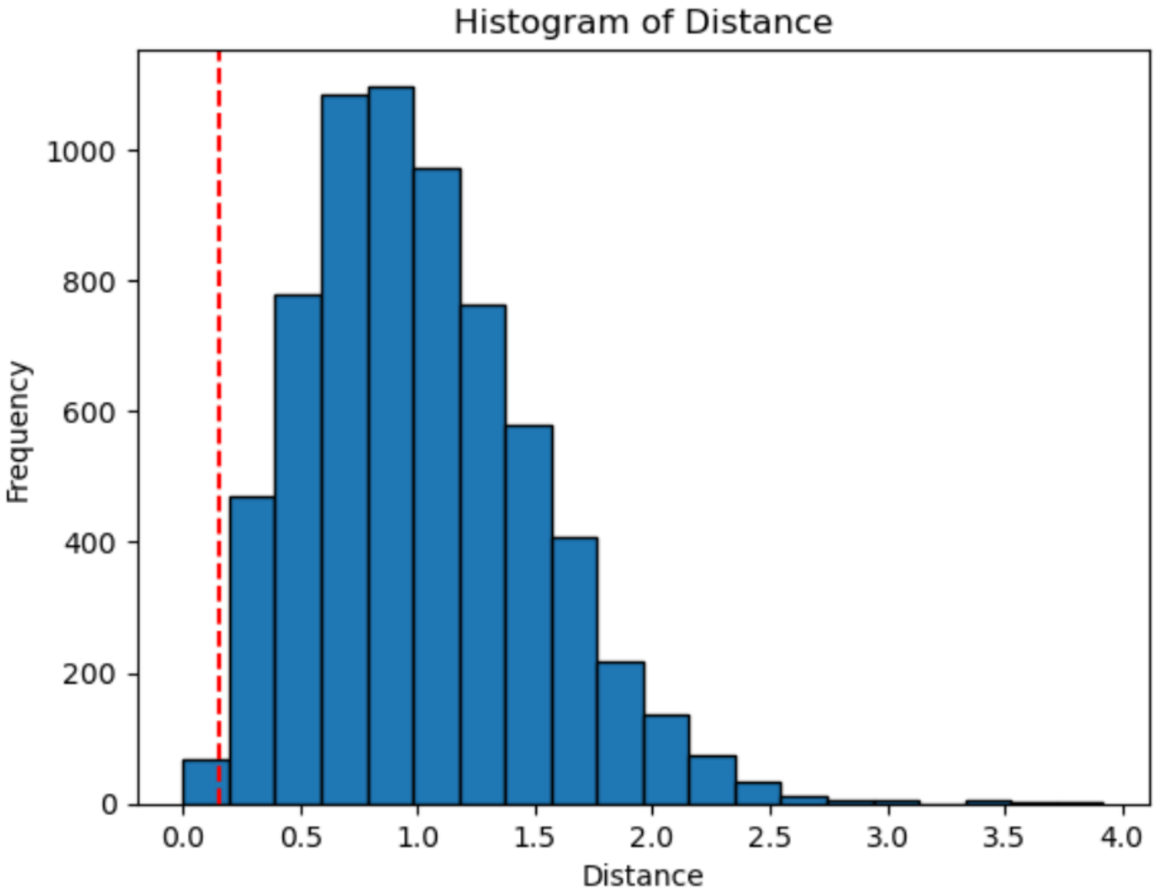

**Fig. S21. Distribution of pairwise haplotype distances.** To be able to distinguish between distinct haplotypes rising to high frequency vs the same haplotype continuing to rise to high frequency in subsequent time periods, we used the Hamming distance metric, defined as the number of positions at which the two sequences differ. We compared the distance between the most common haplotypes with that of the mean distance between randomly selected pairs of haplotypes. Shown is a distribution of distances between randomly chosen haplotypes, normalized by the mean pairwise distance ($d = 20.45$). The red dashed line denotes the 1st percentile (0.15). When the most frequent haplotypes in pairs of time periods had distances below this threshold, they were classified as the same haplotype.

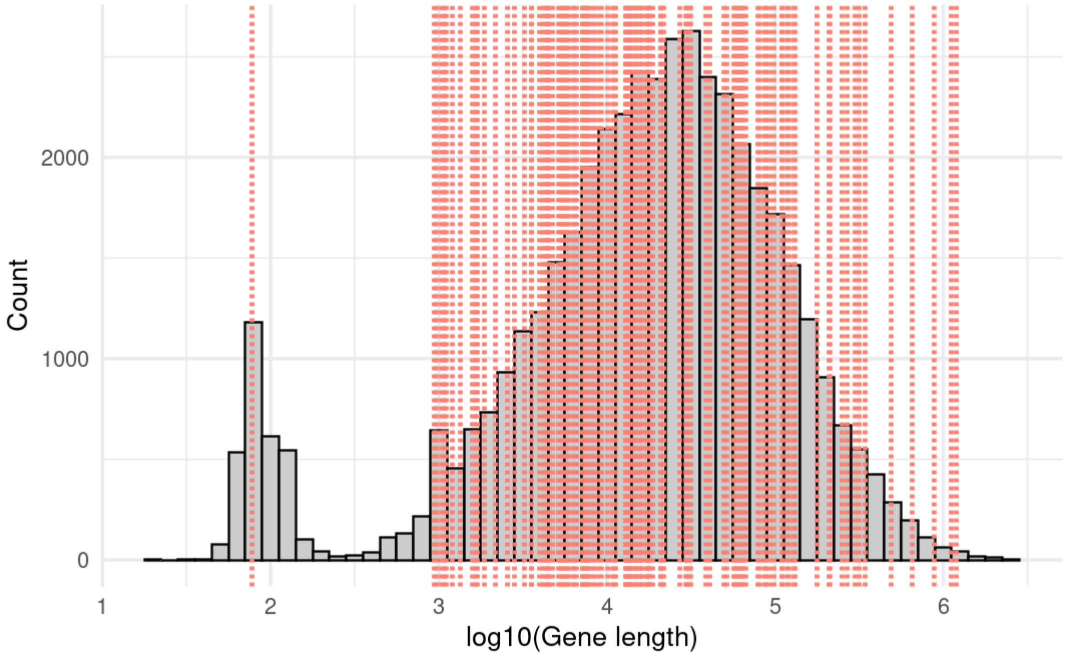

**Figure S22. Length of genes overlapping 14 persistent sweeps.** In grey is the distribution of gene lengths genome wide (23,472 genes). The vertical red lines show the lengths of genes that overlap the 14 sweeps that span across the two ends of the major admixture event that occurred ~4.5 kya (142 genes). The mean length of the genes that overlap sweeps is not significantly longer than expected by chance (p-val =0.326, permutation test comparing the observed mean length to 10,000 random gene sets with same number of genes).

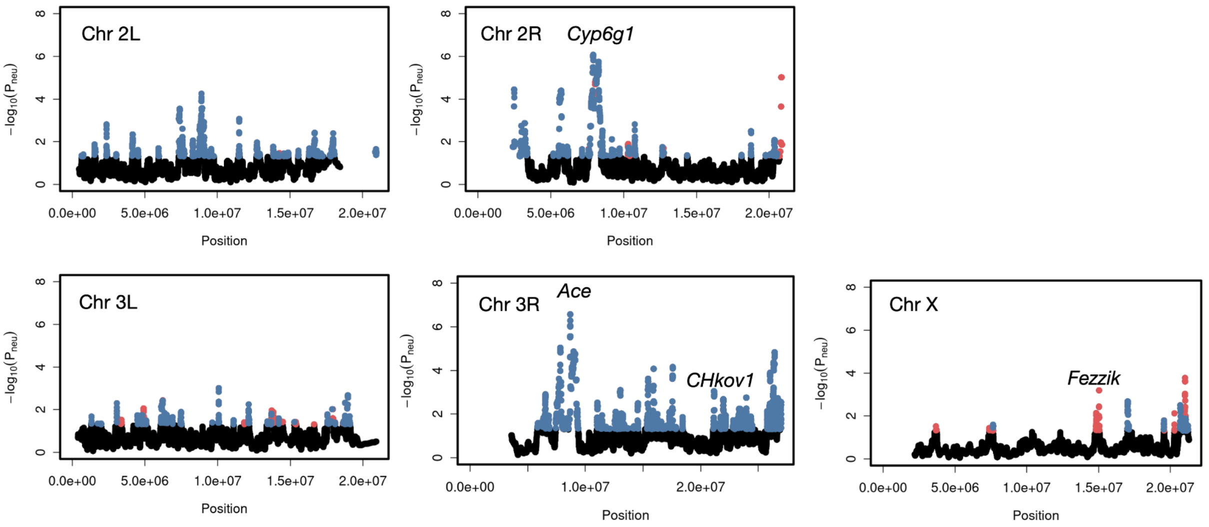

**Fig S23. Genome-wide DANN scan in a North American *D. melanogaster* population.** The y-axis shows the probability of neutrality -log($P_{neu}$) predicted using the DANN. Windows predicted as hard sweeps are colored in red and windows predicted as soft sweeps are colored in blue. Colored in black are windows predicted to be neutral. Previously experimentally validated sweeps reported in the literature are highlighted (*Ace* (1–3) *,* *Cyp6g1(4, 5)*, *CHKov1(6, 7)* , and *Fezzik (8, 9)*).

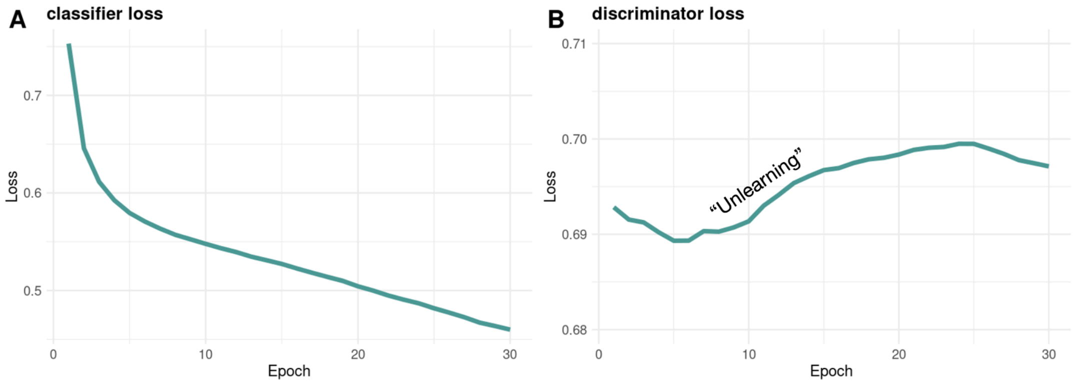

**Figure S24. Classifier and discriminator loss over 30 training epochs.** Shown are example loss curves from training the DANN. **(A)** Categorical cross-entropy loss of the classifier branch gradually decreases over training epochs as the DANN learns to classify neutrality, hard sweeps and soft sweeps. **(B)** Binary cross-entropy loss increases and plateaus as the source and target domains become indistinguishable by the DANN.

### Supplementary tables

| Source | sweeps | hard | soft | unique | 4 time periods | 3 time periods | 2 time periods | 1 time periods |
|---|---|---|---|---|---|---|---|---|
| Constant $N_e$ +43 MD | 127 | 127 | 0 | 71 | 7 | 10 | 15 | 39 |
| Admixture + 43 MD | 91 | 91 | 0 | 48 | 5 | 8 | 12 | 23 |
| Admixture + 5 MD | 101 | 100 | 1 | 57 | 5 | 6 | 17 | 29 |

**Table S2. Number of sweeps detected across four ancient time periods using different source domains for training.** The fist column represents the source domain model that was used to train the DANN. Columns 2-3 show the total number of sweeps, hard sweeps, soft sweeps found across the four time periods. Column 5 shows the total number of unique sweeps across the four time periods. Columns 6-9 show the number of sweeps that show up in all four time periods, in 3 time periods, in two time periods and in only one time period.

| Reference | Sweeps reported | Overlap with DANN | Overlap DANN 1Mb window |
|---|---|---|---|
| Mathieson et al 2015 | 12 | 4 | 5 |
| Pandey, Harris et al, 2024 | 13 | 9 | 9 |
| Le et al. 2022 | 24 | 1 | 4 |
| Akbari et al. 2024 Fig. 2 | 113 | 6 | 14 |
| Kerner et al. 2023 | 89 | 4 | 6 |
| Souilmi et al. 2022 | 57 | 7 | 8 |

**Table S3**. **Overlap between DANN-detected sweeps and those reported in previous studies.** We compare the sweeps identified with the DANN trained with the admixture model with 43% missing data as the source domain to sweeps identified in the six studies listed in this table. In addition, we extended the sweep boundaries identified by the DANN by 1Mb and computed the overlap in this expended window, these results are reported in column 4.

### Supplementary Text

#### Text S1: Ancient DNA data

Data was downloaded from the Allen Ancient DNA Resource (AADR version 51; https://dataverse.harvard.edu/dataset.xhtml?persistentId=doi:10.7910/DVN/FFIDCW).This includes genome-wide data from human populations from Holocene Europe with samples dating from ~7,000 years before present (BP) to ~1,345 BP, covering the Neolithic (N), the Bronze Age (BA), the Iron Age (IA) and the Historical periods (H) (**Fig. 1**).

We focused our analyses on the most reliable samples in our dataset. The criteria we used to select samples are the same as in our previous work (10) and include enrichment for 1240k nuclear targets with an in-solution hybridization capture reagent, removal of samples with high indication of contamination (see (10)), and inclusion of unrelated individuals up to the third degree. Additionally, like in our previous work we selected 177 samples with the highest coverage across all time periods, resulting in a total of 708 aDNA samples (Table S1). Finally, since aDNA coverage is low and thus both alleles at heterozygous sites may not be sampled, and, since there is ascertainment bias of alleles towards the SNPs included in the aDNA capture array (11) we pseudo-haplodized the data by randomly selecting one of the reads that mapped to a given position and assigned that read as the genotype of the sample at that site (10).

#### Text S2: DANN model architecture

The input to the DANN are raw genomic "images" or haplotype matrices (**Fig. 2B**), where rows represent pseudo-haplotypes from each individual and columns represent the ordinal position of each variant site in the sample. These haplotype matrices have dimensions *n* x *L*, where *n*=150 pseudo-haplotypes and *L*=201 SNPs in a given window. In these images, the color of each pixel represents the occurrence of the major or minor allele (12–14). We transformed the alleles into binary values such that the major allele was coded as -1 and minor allele as 1. Missing data was coded as 0. Images were sorted by frequency of most to least frequent haplotype. In text S1 we test the effectiveness of alternative sorting approaches and window sizes.

The feature extractor of the DANN consists of two convolutional layers each containing 64 filters with kernel size of 3x3 and ReLu activation (**Fig. 2A**). Each convolutional layer is followed by 2x2 max-pooling. The last pooling layer is then flattened into a feature vector that is shared between the two subsequent branches of the network: the classifier and the discriminator. Each branch consists of two fully-connected dense layers of 128 neurons each. We use ReLu activation functions and set a dropout rate of 0.5 after each dense layer. The classifier outputs a 3-neuron softmax layer with the predicted probabilities for each of the three classes: hard sweep, soft sweep, or neutral. The discriminator outputs a single sigmoid output layer, which predicts whether the haplotype image comes from the source or target domain.

An important component of the DANN is a GRL between the feature vector and the discriminator branch. During the feed-forwards step of training, the GRL is inactive and the data is passed along to the next layers. During backpropagation, the GRL inverts the gradient of the loss before passing it back to the feature extraction layer (15). This operation penalizes features that discriminate between source and target domains, encouraging the model to learn domain-invariant features critical for accurate classification.

#### Text S3: Haplotype sorting

It has been shown that sorting haplotype images improves the performance of CNNs (12–14, 16). To assess which sorting strategy is most effective for hard and soft sweep classification in the context of aDNA data, we tested three different approaches (**Fig. S1**). The first approach sorts the haplotypes by genomic distance to the most frequent haplotype. By grouping similar haplotypes together, this approach emphasizes haplotype homozygosity and may be well suited for detecting hard sweeps in particular, as a single haplotype rises to high frequency in this scenario, haplotype homozygosity is expected to increase. The second approach ranks the haplotypes from most to least common, capturing patterns where multiple haplotypes are at high frequency, as is expected in the case of soft sweeps. The third approach uses the same frequency-based sorting approach but within a 51 SNP sub-window. Since aDNA data is sparse, genomic windows in the data are very long in terms of base pairs (~450 KB) and thus signatures of selective sweeps may be diluted near the edges of the window. This localized sorting scheme aims to retain the full genomic diversity of the whole window while enhancing sensitivity to signatures of sweeps in a shorter genomic region.

In addition to haplotype sorting, we also tested different window sizes, defined in terms of the number of SNPs. In our previous work (10) we found that window sizes of 201 SNPs, or approximately ~450 KB, were adequate for capturing the signal of selection in this dataset.

However, given the sparse nature of aDNA a 201 SNP-based window can result in very long physical distance, potentially diluting the sweep signal. To address this, we tested a 201 SNP window and a shorter 101 SNP window to evaluate whether reducing the window size improves the performance of our model. Our analysis shows that sorting haplotypes by frequency in 201 SNP windows gives the strongest performance of our method (**Fig. S1**)

#### Text S4: Simulations

We performed simulations of neutrality, hard sweeps and soft sweeps. We use two different demographic models: a constant $N_e = 10^4$ model and a human admixture demographic model that accounts for the major migratory movements contributing to the genetic diversity of contemporary Europeans (17–20)(**Fig. 1C**). In all cases, we simulated a chromosome of length 450kb, reflecting the mean genomic length in aDNA spanning 201 SNPs. We randomly sampled 201 SNPs in the simulated window to match the density and number of SNPs analyzed in the data.

We simulated a total of 400,000 neutral simulations and 400,000 selective sweep simulations (200,000 hard sweeps and 200,000 soft sweeps) under each demographic model. Constant mutation rates and recombination rates were applied to each simulated genome and were drawn uniformly as follows: $\mu \sim U[1e\text{-}8, 1.5e\text{-}8]$ and $\rho \sim U\ [3e\text{-}9, 2e\text{-}8]$. Simulations were performed using SLiM 4.1 (21, 22) followed by recapitation with msprime (23).

We simulated hard sweeps by introducing a single adaptive mutation to the center of the chromosomal segment. We restarted the simulation and re-introduced the adaptive mutation if the mutation was lost. The simulation was allowed to proceed until the adaptive mutation reached a partial frequency (*PF*) drawn from a uniform distribution $PF \sim U[0.5, 0.95]$, for partial sweeps and PF=1.0 for complete sweeps. In all simulations, 177 individuals were sampled and subsequently down sampled to the 150 individuals with the least amount of missing data, reflecting the same procedure applied to aDNA.

We simulated soft sweeps from recurrent *de novo* mutations by introducing adaptive mutations to the center of the chromosome at a rate determined by parameter $\theta_A = 4N_e\mu_A$, where $N_e$ is the effective population size and $\mu_A$ the mutation rate of the adaptive mutation. The value of $\theta_A$ was drawn from a uniform distribution $\theta_A \sim U[1,5]$ as soft sweeps are expected when $\theta_A >= 1$ (24, 25). Finally, the selection strength, *s,* for both hard and soft sweep simulations was drawn from a uniform distribution $s \sim U[0.005, 0.1]$.

Additionally, we simulated sweeps arising from SGV by drawing the frequency of the adaptive mutation prior to the onset of selection from a uniform distribution $f_{init} \sim U[0.025, 0.1]$. Given the computational constraints of simulating a sweep from the SGV jointly with a complex admixture model, we only simulated SGV sweeps using the constant $N_e$ model. Additionally, to improve computational efficiency, we used a hybrid approach where we first simulated a neutral process with msprime and selected a mutation $m_i$ at frequency $f_{init.}$ The resulting tree and mutation id of the variant $m_i$ was then fed to SLiM, where $m_i$ was assigned a selective advantage and allowed to rise in frequency, producing a sweep from SGV.

Simulations were processed to mimic the missing data rate observed in aDNA data. Missing data was added to each site following a beta distribution with a mean of 0.43 per SNP and a standard deviation of 0.28, mimicking the distribution of missing data observed in aDNA (**Fig. S2**). Additionally, we pseudo-haplodized the data by randomly selecting one allele from each sampled individual at each site. Finally, since we ran our scan in 201 SNP windows in aDNA, spanning roughly 450Kbs, we randomly selected 201 SNPs from the simulated chromosome.

#### Text S5: Model training

We implemented the domain-adaptive neural network model using TensorFlow (v2.18.0). All models were trained with the Adam optimizer and a batch size of 64. The classifier branch utilized a categorical cross-entropy loss function, while the discriminator branch used a binary cross-entropy loss function. We fed labeled data from simulations into the classifier branch to compute the class prediction loss ($L_{classifier}$). Simultaneously, we fed a mix of unlabeled data from the source domain (simulations) and target domain (aDNA data) into the discriminator branch to compute the discriminator loss ($L_{discriminator}$). During back propagation, the feature extractor's weights were updated based on a combination of the gradient from the classifier loss and the reversed gradient from the discriminator loss. We use the same simulated data for both branches, however the data is shuffled differently in each mini-batch. The real empirical aDNA data was used on the discriminator branch only. To achieve this training approach, we implement a custom data generator using the Sequence class ('tf.keras.utils.Sequence') that acts as a data generator interface for training Keras models.

The relative contribution of the model's branches can be adjusted via the hyperparameter $\lambda$, such that

$$L_{total} = L_{classifier} + \lambda\ L_{discriminator}$$

If $\lambda$=0 then the GRL is effectively "off" and only the classifier branch learns to classify sweeps without attempting to unlearn domain differences. Conversely, when $\lambda$=1 the GRL is fully "on", assigning equal importance to the tasks of classifying sweeps and unlearning domain differences simultaneously. Following the approach proposed by Ganin & Lempitsky 2014 (15), we gradually increased $\lambda$ from 0 to 1 in order to reduce potential noisy signal from the discriminator at early stages of training. In Ganin & Lempitsky 2014 $\lambda$ is defined as:

$$\lambda = \frac{2}{1 + exp(-\gamma \cdot p)} - 1,$$

Where $p$ is the training progress changing linearly from 0 to 1 and is defined as $p$= $epoch_i / n_{epochs}$, with $n_{epochs}$ =30 total epochs and $i=1,..n_{epochs}$. In all models trained for this paper we set $\gamma$ =10.

We trained the DANN for a total of 30 training epochs. Each epoch took ~282s to train on a single A100 GPU. During training, the discriminator loss gradually increased as the model "unlearned" the misspecification between the source and target. The loss eventually plateaued at ~0.693, consistent with the value expected when the domains become indistinguishable under binary cross-entropy loss. Simultaneously, we observed that the classifier loss monotonically decreased as it learned to distinguish between neutrality, hard sweeps, and soft sweeps (**Fig. S24**).

First, we trained and tested the DANN on simulated source and target domains. In this scenario, we had labeled data from both domains, allowing us to evaluate the performance of the DANN using a validation set from the target domain. In this setting, we selected the model weights from the training epoch that achieved the lowest classifier validation loss when testing on the validation dataset from the target domain.

Next, we trained and tested the DANN but this time using simulated data generated under a human admixture model with an average of 43% missing data per site as the source domain and aDNA data as the target domain. We included all aDNA data for training, where the data only passed through the discriminator branch, not the classifier, since their labels are unknown. The classifier only sees the aDNA data during inference (next section), avoiding any data leakage. In this setting, we could not directly evaluate the performance of the model on a target validation set due to the lack of labeled data. Instead, we assessed performance on simulated data by computing the AUPRC for each class at every training epoch and then averaging across the three classes. We restricted model selection to epochs beyond epoch four, when the GRL begins to influence training ($\lambda$ > 0.5), and chose the epoch with the highest average AUPRC.

#### Text S6: DANN genome-wide scan

We performed a genome-wide scan across all autosomes and all four time periods of aDNA data from this study. We also include a scan on 99 samples of modern humans from the CEU population from the 1000 Genomes Project (26). This modern data scan was generated using a different DANN than the one used on ancient samples, trained on CEU data as the target domain.

To apply the model to all autosomes, we use a sliding window of 201 SNPs, advancing each window by 10 SNPs. To refine the predictions and reduce noise, we then averaged the predicted probabilities every five consecutive, overlapping windows, generating the final predictions used for the scan. The DANN outputs a probability for each of the three classes (neutral, hard sweep or soft sweep). Each window was assigned to the class with the highest probability. To make sure we are identifying distinct selective events, we grouped consecutive non-neutral windows into a single sweep and required sweeps to be at least 1.5 Mb apart. We further demand that consecutive windows have the same class classification within a sweep (with exception is the soft shoulders in HLA and OCA2). Each sweep was represented by the window with lowest probability of neutrality, in other words, the strongest signal within the sweep. The classification of this representative window (hard or soft sweep) was used to classify the whole sweep.

#### Text S7: ABC for hard/soft sweep classification with H12 and H2/H1 statistics

We assessed whether the predictions for hard vs soft sweeps in the data made by the DANN are consistent with predictions made with the haplotype homozygosity statistics, H12 and H2/H1, that can jointly discriminate between hard and soft sweeps (27) . H2/H1 is expected to be small for hard sweeps and large for soft sweeps, conditional on H12 being larger than expected under neutrality. As such, we only analyzed 23 sweeps in total that had an H12 value greater than the 95th percentile of H12 values from 400,000 neutral simulations under the human admixture model as these were least likely to be neutral.

We used an approximate Bayesian computation approach to evaluate if a pair of (H12, H2/H1) values in the data are more likely under a hard vs soft sweep model. To do so, we compare observed values in the data to values measured from simulations of hard sweeps and soft sweeps under the human admixture model. We calculated Bayes factors (BFs) for the observed data by taking the ratio of the number of soft sweep vs hard sweep simulations with a Euclidean distance <0.1 from each (H12, H2/H1) data point (27).

#### Text S8: Jaccard index and permutation test

To quantify the overlap of sweeps across different time periods, we calculated a Jaccard similarity index (J) quantifying sweep overlap between pairs of time periods. J measures the proportion of shared elements between two sets relative to their total combined elements and its defined as

$$J(A,B)=(|A \cap B|)/(|A \cup B|),$$

or in other words, the number of shared elements in sets A and B divided by the total number of elements in A and B. In this scenario A and B represent the sweeps identified in two distinct time periods, such as IA and H. J(IA,H) is obtained by dividing the number of sweeps that are shared between IA and H by the total number of sweeps found across both periods.

Next, to identify whether the value of J was statistically significant, we performed a permutation test where we randomly shuffled the time period labels. We did this 5,000 times generating a null distribution of expected values of J. The p-value of the observed J value is given by the proportion of test statistics from the null distribution that are as extreme or more extreme than the observed value for a pair of time periods.

#### Text S9: Application of a DANN to a North American population of *D. melanogaster*

To assess if the DANN is able to recover known soft sweeps, we applied the DANN to a North American population of *Drosophila melanogaster* that has three well established soft sweeps that have been identified empirically at *Ace (1–3), CHkov1(6, 7)* and *Cyp6g1 (4, 5).* The sweeps at *Ace* and *Cyp6g1* arose from recurrent de novo mutations while the sweep from *CHkov1* arose from standing genetic variation. To run the scan on this data, we trained a new DANN using the *D. melanogaster* data as the target domain and simulated data from a constant $N_e = 10^6$ model with mutation rate $\mu = 1 \times 10^{-9}$ and recombination rate $\rho = 5 \times 10^{-7}$ cM/bp, rescaled by a factor of 50 (28, 29). Our model is able to recover all three known positive controls and classify them as soft sweeps (**Fig. S20**). Additionally we find that soft sweeps dominate across the autosomes of this population and that hard sweeps are enriched on the X chromosome relative to the autosomes, consistent with our previous findings that hemizygosity on the X results in an abundance of hard sweeps (28, 29).

#### References

1. T. Karasov, P. W. Messer, D. A. Petrov, Evidence that Adaptation in Drosophila Is Not Limited by Mutation at Single Sites. *PLOS Genetics* **6**, e1000924 (2010).

2. A. Mutero, M. Pralavorio, J. M. Bride, D. Fournier, Resistance-associated point mutations in insecticide-insensitive acetylcholinesterase. *Proceedings of the National Academy of Sciences* **91**, 5922–5926 (1994).

3. P. Menozzi, M. A. Shi, A. Lougarre, Z. H. Tang, D. Fournier, Mutations of acetylcholinesterase which confer insecticide resistance in Drosophila melanogaster populations. *BMC Evolutionary Biology* **4**, 4 (2004).

4. P. Daborn, S. Boundy, J. Yen, B. Pittendrigh, R. ffrench-Constant, DDT resistance in Drosophila correlates with Cyp6g1 over-expression and confers cross-resistance to the neonicotinoid imidacloprid. *Molecular Genetics and Genomics* **266**, 556–563 (2001).

5. J. M. Schmidt, *et al.*, Copy Number Variation and Transposable Elements Feature in Recent, Ongoing Adaptation at the Cyp6g1 Locus. *PLOS Genetics* **6**, e1000998 (2010).

6. Y. T. Aminetzach, J. M. Macpherson, D. A. Petrov, Pesticide Resistance via Transposition-Mediated Adaptive Gene Truncation in Drosophila. *Science* **309**, 764–767 (2005).

7. M. M. Magwire, F. Bayer, C. L. Webster, C. Cao, F. M. Jiggins, Successive Increases in the Resistance of Drosophila to Viral Infection through a Transposon Insertion Followed by a Duplication. *PLOS Genetics* **7**, e1002337 (2011).

8. A. Glaser-Schmitt, M. J. Wittmann, T. J. S. Ramnarine, J. Parsch, Sexual Antagonism, Temporally Fluctuating Selection, and Variable Dominance Affect a Regulatory Polymorphism in Drosophila melanogaster. *Molecular Biology and Evolution* **38**, 4891–4907 (2021).

9. A. Glaser-Schmitt, J. Parsch, Functional characterization of adaptive variation within a cis-regulatory element influencing Drosophila melanogaster growth. *PLOS Biology* **16**, e2004538 (2018).

10. D. Pandey, M. Harris, N. R. Garud, V. M. Narasimhan, Leveraging ancient DNA to uncover signals of natural selection in Europe lost due to admixture or drift. *Nat Commun* **15**, 9772 (2024).

11. T. Günther, C. Nettelblad, The presence and impact of reference bias on population genomic studies of prehistoric human populations. *PLOS Genetics* **15**, e1008302 (2019).

12. L. Flagel, Y. Brandvain, D. R. Schrider, The Unreasonable Effectiveness of Convolutional Neural Networks in Population Genetic Inference. *Molecular Biology and Evolution* **36**, 220–238 (2019).

13. L. Torada, *et al.*, ImaGene: a convolutional neural network to quantify natural selection from genomic data. *BMC Bioinformatics* **20**, 337 (2019).

14. H. Zhao, N. Alachiotis, Data preprocessing methods for selective sweep detection using convolutional neural networks. *Methods* **233**, 19–29 (2025).

15. Y. Ganin, V. Lempitsky, Unsupervised Domain Adaptation by Backpropagation. [Preprint] (2014). Available at: http://arxiv.org/abs/1409.7495 [Accessed 22 April 2025].

16. R. M. Cecil, L. A. Sugden, On convolutional neural networks for selection inference: Revealing the effect of preprocessing on model learning and the capacity to discover novel patterns. *PLoS Comput Biol* **19**, e1010979 (2023).

17. Y. Souilmi, *et al.*, Admixture has obscured signals of historical hard sweeps in humans. *Nat Ecol Evol* **6**, 2003–2015 (2022).

18. P. D. B. Damgaard, *et al.*, 137 ancient human genomes from across the Eurasian steppes. *Nature* **557**, 369–374 (2018).

19. E. R. Jones, *et al.*, Upper Palaeolithic genomes reveal deep roots of modern Eurasians. *Nat Commun* **6**, 8912 (2015).

20. J. Kamm, J. Terhorst, R. Durbin, Y. S. Song, Efficiently inferring the demographic history of many populations with allele count data. *J. Am. Stat. Assoc.* **115** (2019).

21. B. C. Haller, P. W. Messer, SLiM 4: Multispecies Eco-Evolutionary Modeling. *The American Naturalist* **201**, E127–E139 (2023).

22. B. C. Haller, J. Galloway, J. Kelleher, P. W. Messer, P. L. Ralph, Tree-sequence recording in SLiM opens new horizons for forward-time simulation of whole genomes. *Molecular Ecology Resources* **19**, 552–566 (2019).

23. F. Baumdicker, *et al.*, Efficient ancestry and mutation simulation with msprime 1.0. *Genetics* **220**, iyab229 (2022).

24. P. S. Pennings, J. Hermisson, Soft Sweeps II—Molecular Population Genetics of Adaptation from Recurrent Mutation or Migration. *Molecular Biology and Evolution* **23**, 1076–1084 (2006).

25. P. S. Pennings, J. Hermisson, Soft Sweeps III: The Signature of Positive Selection from Recurrent Mutation. *PLOS Genetics* **2**, e186 (2006).

26. The 1000 Genomes Project Consortium, *et al.*, A global reference for human genetic variation. *Nature* **526**, 68–74 (2015).

27. N. R. Garud, P. W. Messer, E. O. Buzbas, D. A. Petrov, Recent Selective Sweeps in North American Drosophila melanogaster Show Signatures of Soft Sweeps. *PLoS Genet* **11**, e1005004 (2015).

28. M. Harris, N. R. Garud, Enrichment of Hard Sweeps on the X Chromosome in Drosophila melanogaster. *Molecular Biology and Evolution* **40**, msac268 (2023).

29. M. Harris, B. Y. Kim, N. Garud, Enrichment of hard sweeps on the X chromosome compared to autosomes in six *Drosophila* species. *GENETICS* **226**, iyae019 (2024).
